# Tirzepatide reduces diet-induced senescence and NETosis-mediated liver fibrosis in mice

**DOI:** 10.1101/2025.05.27.656284

**Authors:** Feng Chen, QiQi Nam, Liang De Wang, Felicia Chammas, Torsten Wuestefeld, Bernett Teck Kwong Lee, Siu Ling Wong

## Abstract

Lipofuscin, a protein-lipid complex that progressively accumulates in senescent cells, was commonly observed in the liver biopsies from patients with liver fibrosis or cirrhosis regardless of the etiological insults. However, whether and how lipofuscin contributes to the development of liver fibrosis remains unknown. Using a mouse model of liver fibrosis induced by choline-deficient L-amino acid-defined high-fat diet (HFD), we found that lipofuscin accumulation preceded the emergence of liver fibrosis. A significant, positive and strong correlation was observed between the hepatic lipofuscin levels and liver fibrosis severity. Neutrophils were recruited to the liver in mice fed HFD, and those that were in close proximity to lipofuscin-laden hepatocytes displayed higher susceptibility to produce neutrophil extracellular traps (NETs). We revealed that lipofuscin-laden hepatocytes generated CXCL5, which promoted NETosis. When mice defective in NETosis (*Vav1-Cre Padi4^fl/fl^*) were fed HFD, they had significantly less collagen deposition in the liver, indicating the critical role of NETs in the development of liver fibrosis associated with lipofuscin accumulation. Notably, we showed that tirzepatide, the latest glucagon-like peptide-1 receptor and glucose-dependent insulinotropic polypeptide receptor dual agonist recently approved for type 2 diabetes, attenuated NETosis-dependent liver fibrosis by preventing hepatic lipofuscin accumulation, and reduced other hepatic senescence signatures as well. In summary, our study unveiled the pathogenic, NETosis-enhancing effect of lipofuscin in liver fibrosis, and demonstrated the potential of repurposing tirzepatide for combating liver fibrosis associated with lipofuscin and for preventing organ senescence.

## Introduction

Chronic liver disease (CLD) and the associated complications account for approximately 1 million deaths worldwide annually^1^. The most common type of CLD is metabolic dysfunction-associated steatotic liver disease (MASLD)^2^, amongst which lean MASLD is highly prevalent in Asia^3^ and is associated with a higher risk of all-cause mortality compared to obese MASLD^4^. CLD, including MASLD, encompasses a spectrum of progressive disorders in the liver, ranging from steatohepatitis, fibrosis, to cirrhosis and potentially hepatocellular carcinoma. In the CLD spectrum, fibrosis is the last stage that is reversible^5^. Upon progression to cirrhosis, patients have a 5- to 10-times higher mortality risk than the general population^6^. Of note, cirrhosis is the most important risk factor for hepatocellular carcinoma which ranks the third amongst cancer-related deaths globally^7^. While patients with CLD may still reduce the severity of steatosis with lifestyle management, treatment and approved medications for liver fibrosis remain a significant unmet need clinically. Mitigating pathogenesis at fibrosis is therefore crucial for preventing progression to cirrhosis and liver failure.

Lipofuscin is commonly observed in human liver biopsies from patients with CLD^8,9^, which can result from various insults such as high-fat diet^10^, drug^11^, and viral infection^5^. Lipofuscin is an intracellular pigment mainly composed of oxidized proteins and lipids, with small amounts of carbohydrates and metals^12^. As one of the hallmarks of cellular senescence, lipofuscin also forms in the liver of aged humans^13^ and animals^14^. Lipofuscin has long been regarded as an inert byproduct of aging without much pathological impact. Until recently, studies began to suggest that lipofuscin can facilitate disease progression^15,16^, such as promoting retinal degeneration^15^. However, whether and how lipofuscin is involved in the development of liver fibrosis remains unknown.

Immune activation is involved in organ inflammation, healing, and even over-healing, which can give rise to fibrotic scarring response. When the liver is under stress or injured, neutrophils are one of the main immune cells that are rapidly recruited to the liver. Instead of tissue repair, hyperactive neutrophils can contribute to the development and progression of liver disease^17,18^. They can release neutrophil extracellular traps (NETs) that consist of decondensed chromatin decorated with cytotoxic granule proteins^19^. First discovered to kill bacteria for controlling infection^19,20^, NETs also form in sterile inflammation and fuel diseases such as diabetic complications^21–25^ and organ fibrosis^26,27^. Thus far, whether NET formation mediates liver disease associated with lipofuscin and how they play a role in liver fibrosis remains to be elucidated.

While there are limited treatment options for liver fibrosis, a recent clinical trial showed in the explorative secondary endpoint that tirzepatide (TZP), a glucagon-like peptide-1 receptor (GLP1R) and glucose-dependent insulinotropic polypeptide receptor (GIPR) dual agonist that was approved for the treatment of type 2 diabetes in 2022^28^, reduces liver fibrosis in patients with metabolic dysfunction-associated steatohepatitis (MASH)^29^. The mechanism underpinning the anti-fibrotic effect of TZP and whether TZP is effective for liver fibrosis with hepatic lipofuscin accumulation which is common in CLD arisen from different causes warrants investigation.

In this study, we sought to explore whether and how lipofuscin contributed to liver fibrosis, and to examine if TZP could alleviate liver fibrosis by intercepting lipofuscin accumulation and neutrophil activation. We adopted a choline-deficient L-amino acid-defined high-fat diet (CDAHFD, later abbreviated as HFD)-induced mouse model of liver fibrosis without weight gain^30^, potentially representing patients with lean MASLD. Choline deficiency in this model also resembles decreased phosphatidylcholine (product of choline) in MASH^31^. Importantly, this HFD promotes robust lipofuscin deposition in the liver^8^, which could be attributed to oxidative stress^32,33^. We found that hepatic lipofuscin accumulated before the development of liver fibrosis in this model. Lipofuscin-containing hepatocytes produced CXCL5, predisposing neutrophils to undergo NETosis that consequentially led to liver fibrosis. Finally, administration of TZP decreased NETosis-dependent liver fibrosis by reducing levels of lipofuscin in the liver, and normalized other senescence biomarkers in the liver of mice fed the HFD.

## Results

### Accumulation of lipofuscin precedes collagen deposition in the liver

Yellowish-brown pigment was observed in the hematoxylin and eosin (H&E)-stained liver sections from mice fed HFD for 10 weeks (Fig. 1a). When unstained sections were observed under the fluorescence microscope, the pigments appeared autofluorescent (Fig. 1a). In contrast, no yellowish-brown pigments nor autofluorescent substance was observed in the liver sections from mice fed control diet (CD) for the same duration (Fig. 1a). Sudan Black B (SBB) staining^34^ confirmed that the autofluorescent substances were lipofuscin (Fig. 1b). Immunostaining of asialoglycoprotein receptor 1 (ASGR1, a membrane protein specific to hepatocytes) and lysosomal-associated membrane protein 1 revealed that lipofuscin (black patches observed in the differential interference contrast channel) accumulated in the hepatocytes and was contained within the lysosome (Fig. 1c,d). This observation aligned with previous studies on other tissues, such as the retinal pigment epithelium and the heart, that lipofuscin was found in the lysosomal compartment of the cells^16,35^.

**Fig. 1.**
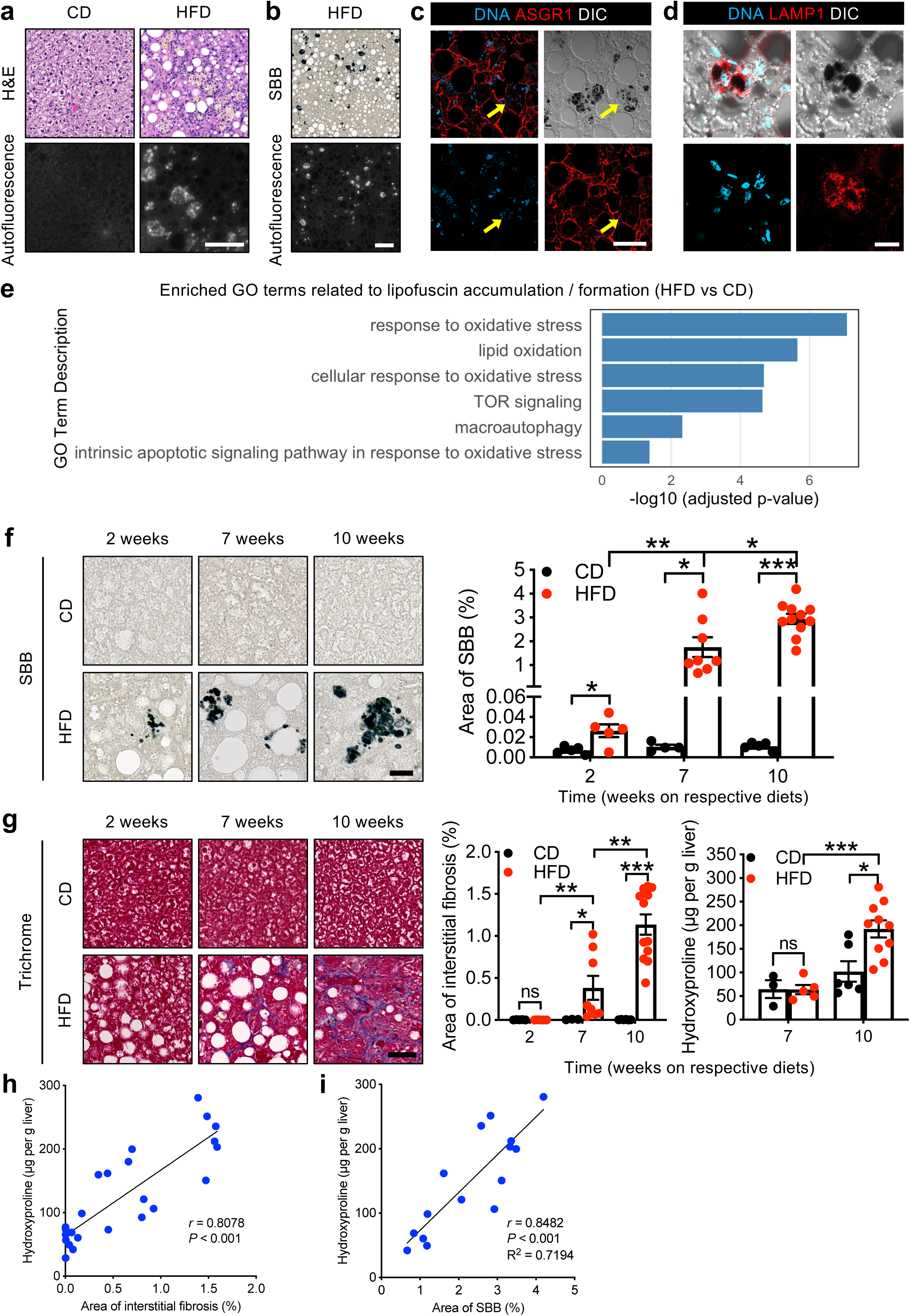
Lipofuscin accumulation in the liver precedes liver fibrosis, and it significantly and positively correlates with the severity of liver fibrosis. **a-e,** Male C57BL/6J mice were fed control diet (CD) or high-fat diet (HFD) for 10 weeks. **a,b,** Images of hematoxylin and eosin (H&E) staining (**a**), autofluorescence imaging (**a,b**), and Sudan Black B (SBB) staining (**b**) on formalin-fixed paraffin-embedded (FFPE) liver sections from mice. Scale bar, 100 µm. **c,d,** Immunofluorescence staining for hepatocyte membrane marker, asialoglycoprotein receptor 1 (ASGR1) (**c**) or lysosomal-associated membrane protein 1 (LAMP1) (**d**) on the liver sections. Dark patches shown in the differential interference contrast (DIC) channel were SBB-stained lipofuscin. Yellow arrows indicate lipofuscin-containing cells. Single-plane images were acquired for slides stained for ASGR1 (**c**). Maximum intensity projection images were generated from Z-stack images for slides stained for LAMP1 (**d**). Scale bars, 50 µm. **e**, Gene Ontology (GO) terms related to lipofuscin accumulation and formation enriched in the liver of mice fed HFD compared to mice fed CD for 10 weeks. **f**, Representative images of lipofuscin stained by SBB (left) and quantification of area of SBB (%) (right) on FFPE liver sections from mice fed the respective diets for 2, 7, and 10 weeks. Left to right on the bar chart, *n* = 5, 5, 4, 8, 5, 11. Scale bar, 20 µm. **g**, Representative images of liver interstitial fibrosis stained by Masson’s trichrome (left) and quantification of area of interstitial fibrosis (%) (middle) on FFPE liver sections from mice fed the respective diets for 2, 7, and 10 weeks. Level of liver fibrosis assessed by hydroxyproline content (µg per g liver) in mice fed the respective diets for 7 and 10 weeks (right). For quantification of area of interstitial fibrosis (%), left to right on the bar chart, *n* = 5, 5, 3, 8, 6, 12. For hydroxyproline assay, left to right on the bar chart, *n* = 3, 5, 6, 10. Scale bar, 50 µm. Statistical comparisons were made using two-tailed unpaired t-test (**f**) or Mann-Whitney test (**g**). \**P* < 0.05, \*\**P* < 0.01 and \*\*\**P* < 0.001. Data are mean ± s.e.m. **h**, Spearman correlation analysis conducted between area of interstitial fibrosis (%) revealed by Masson’s trichrome staining and hepatic hydroxyproline content (µg per g liver) from mice fed CD or HFD for 7 and 10 weeks. *n* = 3 for 7-week CD group, *n* = 5 for 7-week HFD group, *n* = 6 for 10-week CD group, *n* = 10 for 10-week HFD group. **i**, Pearson correlation analysis performed between area of SBB (%) and hepatic hydroxyproline content (µg per g liver) from mice fed HFD for 7 and 10 weeks. *n* = 5 for 7-week HFD group, *n* = 10 for 10-week HFD group.

Bulk RNA sequencing (RNA-seq) also unveiled that HFD induced a milieu facilitating lipofuscin accumulation in the liver. Due to its heterogeneous composition^36^, lipofuscin lacks specific RNA signatures. Hence, we examined changes in pathways that could implicate lipofuscin formation or accumulation^37^. Livers from HFD-fed mice had a significant enrichment in biological pathways of oxidative stress and autophagy (Fig. 1e), both of which have been shown to be involved in lipofuscin formation^32,38^.

To delineate the timeframe of lipofuscin emergence and liver fibrosis development, mice were fed shorter durations of the HFD. We observed that lipofuscin started to accumulate in the liver of the mice after being on HFD for 2 weeks, and the amount increased progressively across 10 weeks, whereas CD-fed mice did not show lipofuscin accumulation at all (Fig. 1f). Interestingly, pathological collagen deposition did not take place until 7 weeks of HFD, by then the liver fibrosis severity was still relatively mild, as the increase in hepatic collagen was only observable using Masson’s trichrome staining but not yet detected by the less sensitive hydroxyproline assay (Fig. 1g). Liver fibrosis was detected using both methods when mice were fed 10 weeks of HFD (Fig. 1g). Other pathological features such as hepatocyte ballooning, immune cell infiltration and steatosis were observed at 2 weeks post HFD and onwards (Extended Data Fig. 1a). Quantification of Oil Red O staining reflected a slight reduction in steatosis in mice fed HFD for 10 weeks compared to those on HFD for 7 weeks (Extended Data Fig. 1b), which could be attributed to the development of fibrosis^39^.

### Hepatic lipofuscin levels significantly and positively correlate with severity of liver fibrosis

The relationship between lipofuscin levels and severity of the pathological features in HFD-induced CLD was examined. H&E-stained liver sections were graded according to the steatosis, activity (including hepatocyte ballooning and immune cell infiltration), and fibrosis (SAF) score to assess liver injury broadly, a method clinically adopted to evaluate liver damage in patients^40^. Amount of hepatic lipofuscin positively and significantly correlated with SAF score (*r* = 0.5964, *P* = 0.02) (Extended Data Fig. 1c), and the predictive power of hepatic lipofuscin was the most pronounced for the fibrosis parameter (R^2^ = 0.6223, *P* < 0.001) (Extended Data Fig. 1d-f). Such finding was recapitulated in the correlation analysis using more quantitative measurements of fibrosis. In fact, area of interstitial fibrosis quantified from trichrome-stained liver sections correlated well with hydroxyproline levels of liver homogenates (*r* = 0.8078, *P* < 0.001, Fig. 1h), suggesting that both measurements were reliable for quantifying the extent of liver fibrosis. We found that lipofuscin levels strongly correlated with hepatic hydroxyproline levels (*r* = 0.8482, *P* < 0.001) and were highly predictive of severity of liver fibrosis (R^2^ = 0.7194) (Fig. 1i).

### Neutrophils in the liver but not in the blood are predisposed to NETosis in HFD-fed mice

To explore potential pathways that underpinned HFD-induced liver fibrosis, bulk RNA-seq analysis was performed on the liver tissue of mice fed CD or HFD for 10 weeks. Volcano plot showed 6,072 differentially expressed genes (DEGs) with known biological functions; amongst which, 3,467 DEGs were significantly upregulated while 2,605 were significantly reduced in the liver of HFD-fed mice compared to that of mice fed CD (Fig. 2a). In addition to metabolic pathways, there was also significant enrichment in biological processes involving immune regulation, such as leukocyte migration, chemotaxis and myeloid leukocyte activation (Fig. 2b). Since neutrophils have been implicated in organ fibrosis^26,27^ and progression of CLD^17,18^, we examined changes in neutrophil-related Gene Ontology (GO) terms. Neutrophil migration, chemotaxis and activation pathways were found to be significantly enriched in the liver of the HFD-fed mice (Fig. 2c). The bulk RNA-seq findings suggested the critical involvement of neutrophils and its activation in liver fibrosis featured with lipofuscin accumulation.

**Fig. 2.**
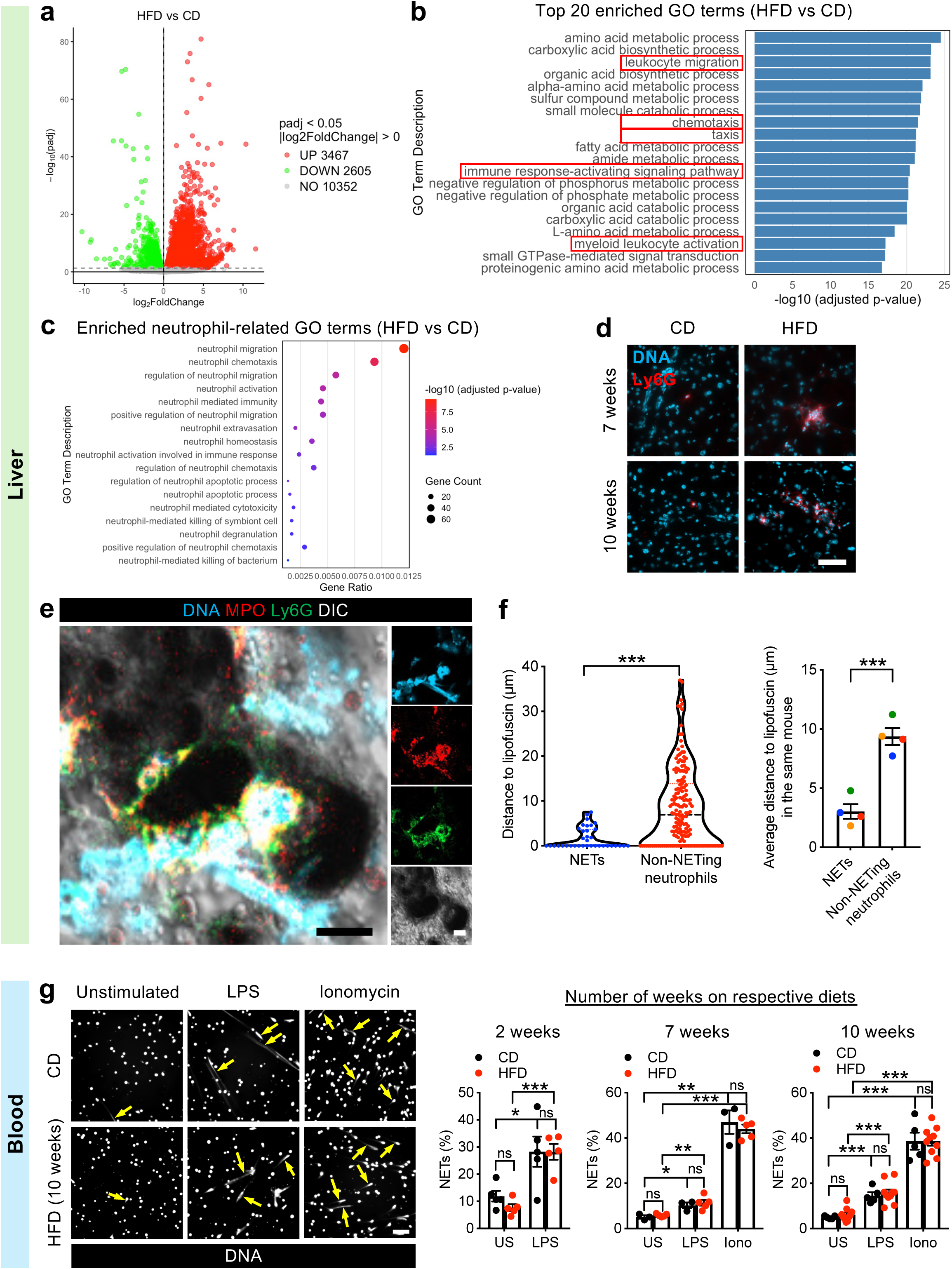
Neutrophils in the liver but not in the blood are more prone to NETosis in HFD-fed mice. **a-c,** Bulk RNA sequencing (RNA-seq) was performed on the liver of mice fed CD or HFD for 10 weeks (*n* = 3 per group). **a,** Volcano plot of differentially expressed genes (DEGs) significantly upregulated (red) or downregulated (green) in the liver. **b,c,** Top 20 enriched GO terms based on adjusted p-value (**b**) and enriched neutrophil-related GO terms (**c**) in the liver. **d,** Immunofluorescence staining for neutrophil recruitment (Ly6G^+^ cells) in the liver of mice fed HFD or CD for up to 10 weeks. Scale bar, 50 µm. **e,** Single-plane confocal images showing NETs [externalized DNA co-localized with myeloperoxidase (MPO) from Ly6G^+^ cells] in the liver of mice fed HFD for 10 weeks. DIC channel showed SBB-stained lipofuscin (black patches). Scale bars, 5 µm. **f,** The shortest distance between NETs or non-NETing neutrophils to lipofuscin (µm) measured in the liver sections from mice fed HFD for 10 weeks. Each point represents an individual neutrophil (*n* = 168) or NET (*n* = 36) (left) or average distance of NETs or non-NETing neutrophils to lipofuscin (µm) calculated per mouse (*n* = 4) (right). Statistical comparisons were made using Mann-Whitney test (left) or two-tailed unpaired t-test (right). Data are expressed as median with a quartile (left) or mean ± s.e.m. (right). **g,** Blood neutrophils were isolated from mice fed the respective diets for up to 10 weeks and stimulated with 10 µg/mL lipopolysaccharide (LPS) or 4 µM ionomycin (Iono) for 4 hours. Representative images of NETs produced by neutrophils after *ex vivo* stimulation (left) and quantification of NETs (%) from the isolated blood neutrophils (right). Yellow arrows indicate examples of NETs. US, unstimulated. *n* = 5 per group for 2-week diet, *n* = 3 for 7-week CD group, *n* = 5 for 7-week HFD group, *n* = 5 for 10-week CD group, *n* = 10 for 10-week HFD group. Scale bar, 100 µm. Statistical comparisons were made using two-tailed unpaired t-test. \**P* < 0.05, \*\**P* < 0.01, \*\*\**P* < 0.001, ns, not significant. Data are mean ± s.e.m.

Immunostaining of the liver sections revealed similar findings as bulk RNA-seq. An increase in the recruitment of neutrophils (Ly6G^+^ cells) was observed in the liver of mice fed HFD for 7 and 10 weeks (Fig. 2d). Interestingly, NETs (externalized DNA colocalized with myeloperoxidase) were generated by neutrophils that were in close proximity to the lipofuscin-laden hepatocytes (Fig. 2e). The distance between NETs and lipofuscin was remarkably shorter than that between non-NETing neutrophils and lipofuscin (Fig. 2f), suggesting that neutrophils close to lipofuscin-laden cells were more prone to NETosis, while those that were further away were not as susceptible. Surprisingly, NETosis propensity of blood neutrophils was the same between CD- and HFD-fed mice when these cells were stimulated *ex vivo* with known NETosis stimulants, regardless of the diet duration (Fig. 2g). Collectively, these results suggested that the liver microenvironment is essential for promoting NETosis in mice fed the HFD.

### CXCL5 generated by lipofuscin-laden hepatocytes enhances NET formation

DEGs of the top 20 GO terms that were related to immune regulation were further examined; one-third of them were classified under cytokine/chemokine-related pathways (Extended Data Fig. 2). GO enrichment analysis showed that biological processes of cytokine and chemokine production were indeed enriched in the liver of HFD-fed mice (Fig. 3a). Compared to those on CD, the cytokine/chemokine DEG that exhibited the greatest fold change (log2foldchange ∼ 8.4) and a statistically significant upregulation in the liver of HFD-fed mice was *Cxcl5* (Fig. 3b). CXCL5, encoded by *Cxcl5*, is a chemoattractant that facilitates neutrophil migration and activates neutrophils by inducing reactive oxygen species generation^41,42^. Immunostaining of liver sections from HFD-fed mice showed that CXCL5 was preferentially expressed in lipofuscin-containing cells, around which neutrophils were often clustered (Fig. 3c).

**Fig. 3.**
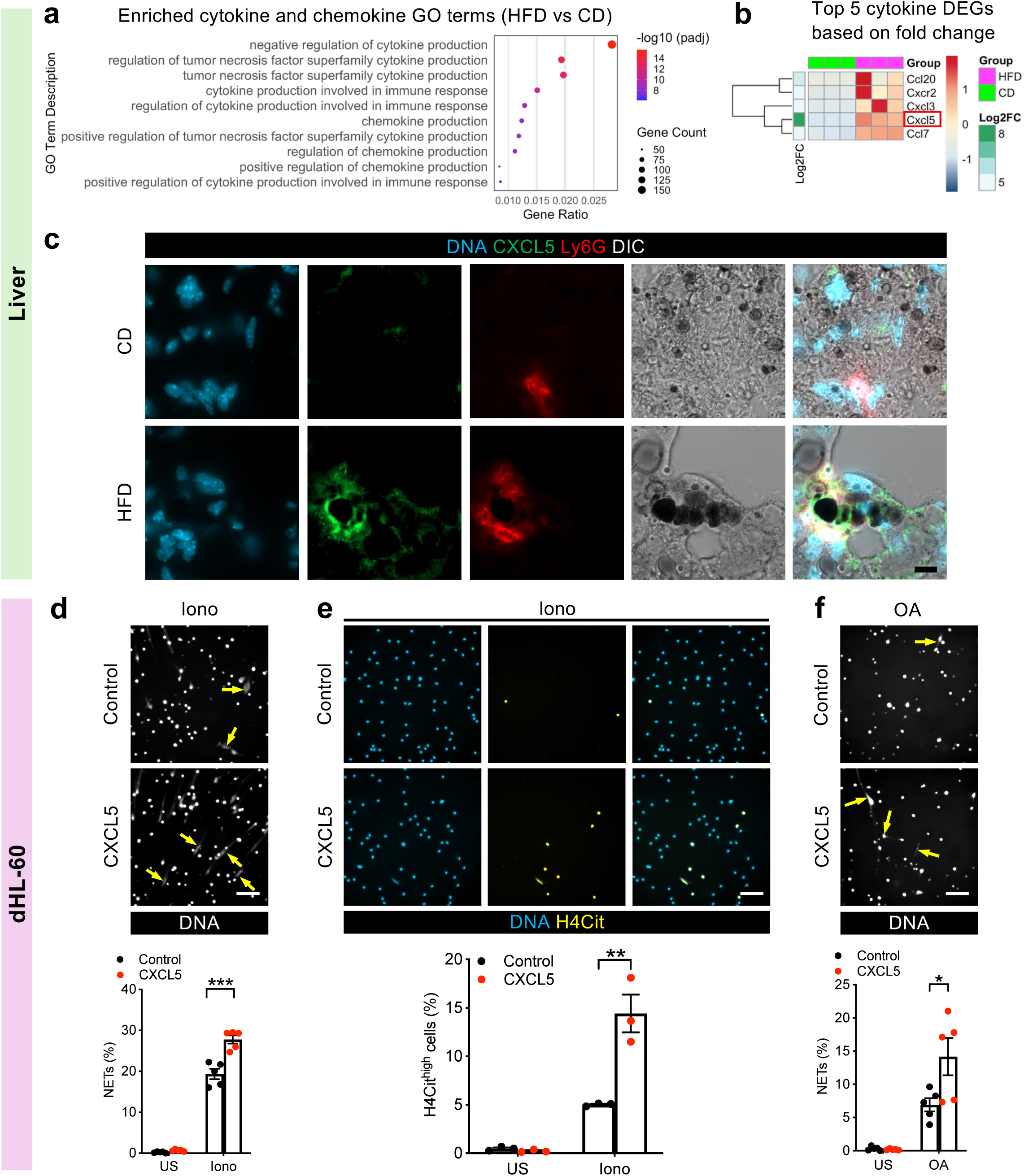
CXCL5 expressed by lipofuscin-laden hepatocytes promotes NETosis. **a,b,** Bulk RNA-seq was performed on the liver of mice fed CD or HFD for 10 weeks (*n* = 3 per group). Top 10 enriched cytokine and chemokine-related GO terms (**a**) and heatmap of top 5 cytokine/chemokine DEGs based on fold change (**b**) in the liver of mice on HFD compared to mice on CD for 10 weeks. padj, adjusted p-value. Log2FC, log2FoldChange. **c,** Images of immunofluorescence staining of CXCL5 and Ly6G on the liver of mice fed HFD or CD for 10 weeks. DIC channel showed SBB-stained lipofuscin (black patches). Scale bar, 20 µm. **d,e,** dHL-60 were incubated with 50 ng/mL CXCL5 for 24 hours and stimulated with 4 µM ionomycin for 4 hours (**d**) or 1 hour (**e**). **d,** Representative images of NETs from dHL-60 (top) and quantification of NETs (%) (below). Yellow arrows indicate examples of NETs. *n* = 5 per group. Scale bar, 100 µm. **e,** Representative images of H4Cit^high^ cells from dHL-60 (top) and quantification of H4Cit^high^ cells (%) (bottom). *n* = 3 per group. Scale bar, 100 µm. **f,** dHL-60 were incubated with 50 ng/mL CXCL5 for 24 hours and stimulated with 50 µM oleic acid (OA) for 4 hours. Representative images of NETs (top) and quantification of NETs (%) (bottom). Yellow arrows indicate examples of NETs. *n* = 5 per group. Scale bar, 100 µm. US, unstimulated. Iono, ionomycin. Statistical comparisons were made using two-tailed unpaired t-test. \**P* < 0.05, \*\**P* < 0.01 and \*\*\**P* < 0.001. Data are mean ± s.e.m.

To examine if CXCL5 predisposed neutrophils for NETosis, we performed *in vitro* cellular experiments using human HL-60-differentiated neutrophils (dHL-60 cells), which are widely adopted for studying neutrophil activation and NETosis^43–45^. The rationale of using dHL-60 cells in place of blood neutrophils is also due to the limitation that blood neutrophils are too short-lived for *ex vivo* experimentation. dHL-60 cells were incubated with recombinant human CXCL5 overnight before being stimulated for NETosis. While CXCL5 alone did not trigger NETosis, 44% more neutrophils with CXCL5 exposure produced NETs when stimulated by ionomycin, compared to neutrophils without CXCL5 exposure (Fig. 3d). Peptidylarginine deiminase 4 (PAD4) is a crucial enzyme for NETosis, where it decondenses the chromatin by changing the positively-charged arginine residues on histones to neutral citrulline^46^. Level of citrullinated histones is therefore adopted as a marker for PAD4 activity. We observed that there were concomitantly more H4Cit^high^ neutrophils after the cells were exposed to CXCL5, indicating that CXCL5 promoted NETosis through PAD4 (Fig. 3e). The NETosis experiment was also repeated using a stimulant that better simulated the liver microenvironment in this mouse model. Increased levels of fatty acids were reported in MASH, one of which was oleic acid^47^. Stimulating dHL-60 cells with oleic acid alone triggered ∼ 7% of neutrophils to undergo NETosis and the number increased to 14% when the neutrophils were pre-incubated with CXCL5 (Fig. 3f). In summary, CXCL5 generated by lipofuscin-laden hepatocytes possibly promotes NETosis in the inflamed liver microenvironment.

### NETs mediate liver fibrosis that is presented with lipofuscin

Since NETs were observed in the liver of mice fed with HFD, we sought to understand if NETs play a role in the development of liver fibrosis. Hematopoietic cell-specific PAD4-deficient mice (*Vav1-Cre Padi4^fl/fl^*, later abbreviated as KO), which were not able to produce NETs^48^, were fed CD or HFD for 10 weeks (Fig. 4a). Consistent with our findings using wild-type mice (Fig 2e,f), neutrophils close to the lipofuscin-laden cells formed NETs in the liver of HFD-fed *Padi4^fl/fl^* (control counterparts for the KO) (Fig. 4b). In contrast, neutrophils of HFD-fed KO mice appeared intact with myeloperoxidase localized in the cytosol and did not release NETs despite their close distance to lipofuscin-laden cells (Fig. 4b). Notably, less liver fibrosis was observed in HFD-fed KO mice in comparison to the control mice (Fig. 4c), indicating a critical role of NETs in liver fibrosis. Surprisingly, no reduction in the amount of hepatic lipofuscin was observed in the KO mice on HFD when compared to the control counterparts (Fig. 4d), and neither was in the hepatic levels of CXCL5 (Fig. 4e). These findings suggested that lipofuscin accumulation and CXCL5 production were not NETosis-dependent, and confirmed that they were both upstream of NET formation in the liver. There were also no noticeable differences in steatosis, hepatocyte ballooning and immune cell recruitment between control and KO mice when both were fed HFD (Fig. 4f,g). Collectively, these results showed that NETosis is involved in lipofuscin-associated liver fibrosis. Intercepting NET formation could reduce liver fibrosis, although lipofuscin accumulation and increase in CXCL5 are not corrected.

**Fig. 4.**
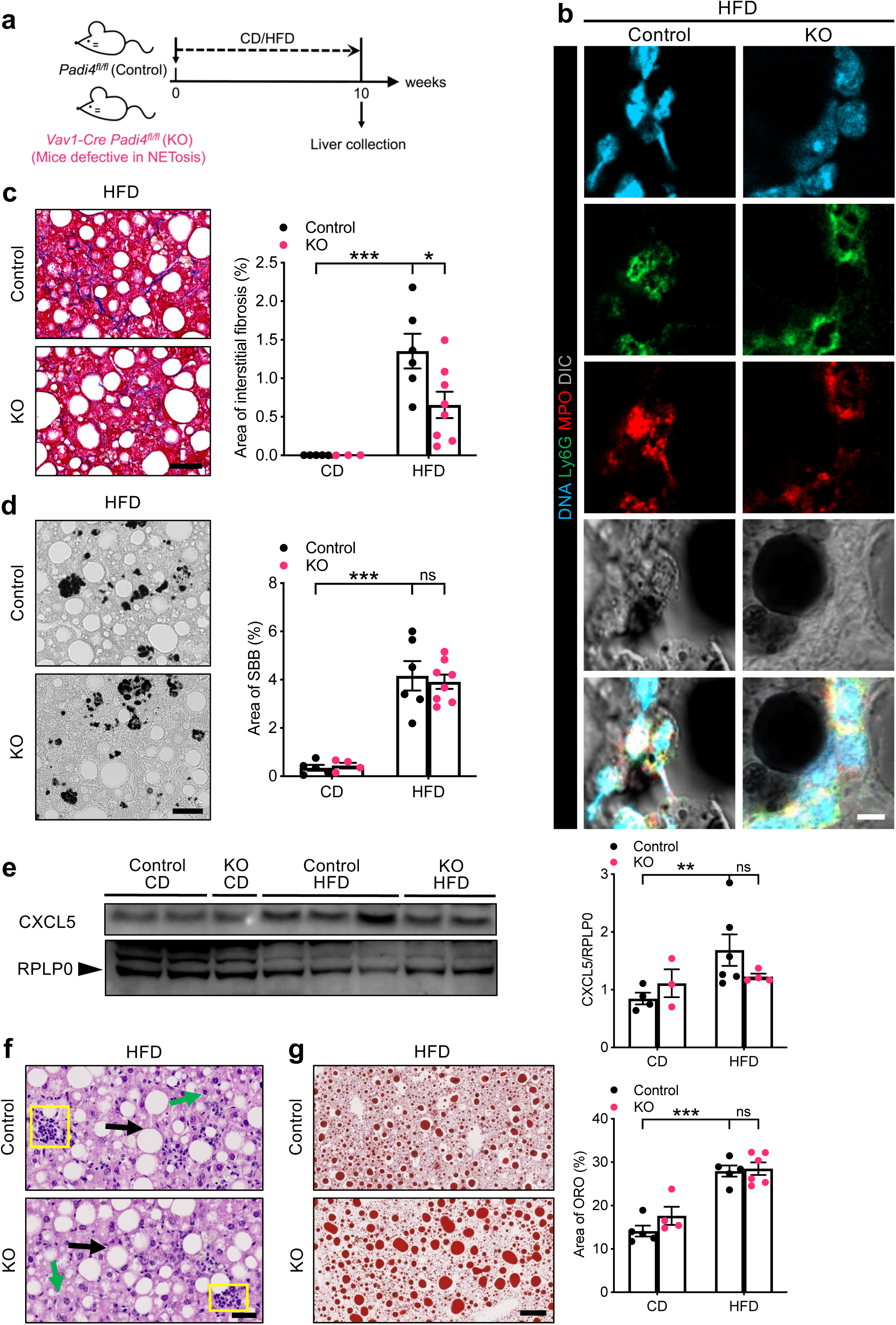
Liver fibrosis is decreased in mice defective in NETosis without reducing lipofuscin accumulation. **a-g,** Male *Padi4^fl/fl^*(Control) and *Vav1-Cre Padi4^fl/fl^* (KO) fed CD or HFD for 10 weeks (**a**). **b,** Single-plane confocal images of neutrophils and NETs in the liver of control and KO mice fed HFD for 10 weeks. DIC channel showed SBB-stained lipofuscin (black patches). Scale bar, 5 µm. **c,** Representative images of liver interstitial fibrosis revealed by Masson’s trichrome (left) and quantification of area of interstitial fibrosis (%) (right) in the liver sections. Left to right on the bar chart, *n* = 5, 3, 6, 8. Scale bar, 50 µm. **d,** Representative images of lipofuscin stained by SBB (left) and quantification of area of SBB (%) (right) in the liver. Left to right on the bar chart, *n* = 5, 4, 6, 8. Scale bar, 50 µm. **e,** Representative Western blots (left) and quantification (right) of CXCL5 levels in the liver. Ribosomal protein lateral stalk subunit P0 (RPLP0) served as the loading control. Left to right on the bar chart, *n* = 4, 3, 6, 4. **f,** Representative images of H&E-stained liver sections from control and KO mice on HFD for 10 weeks, showing hepatocyte ballooning (green arrows), steatosis (black arrows) and immune cell infiltration (yellow squares). Scale bar, 50 µm. **g,** Representative images of ORO-stained liver sections (left) and quantification of area of ORO (%) (right). Left to right on the bar chart, *n* = 5, 4, 5, 6. Scale bar, 50 µm. Statistical comparisons were made using two-tailed unpaired t-test (**c,d,g**) or Mann-Whitney test (**e**). \**P* < 0.05, \*\**P* < 0.01, \*\*\**P* < 0.001, ns, not significant. Data are mean ± s.e.m.

### TZP diminishes liver fibrosis by reducing hepatic lipofuscin accumulation

Since TZP was shown to reduce liver fibrosis in patients with obesity^29^, we aimed to examine if TZP could inhibit the NETosis pathway and hence reduce liver fibrosis that involved lipofuscin. We employed two treatment regimens to examine the effect of TZP on lipofuscin-associated liver fibrosis. First, concurrent treatment, in which TZP (or vehicle) treatment started at the same time as the diets for 10 weeks (Extended Data Fig. 3a), allowed us to evaluate whether TZP could prevent the onset of liver fibrosis; second, delayed treatment, in which mice were fed CD or HFD for 7 weeks, followed by TZP (or vehicle) treatment for an additional 10 weeks along with the respective diets (Fig. 5a), better mimicked the clinical scenario where patients sought medical attention when symptoms of liver fibrosis became apparent. Both TZP treatment regimens drastically lessened liver fibrosis in HFD-fed mice, as reflected by Masson’s trichrome staining and hydroxyproline assay (Extended Data Fig. 3b and Fig. 5b). Unexpectedly, both regimens of TZP treatment remarkably reduced the amount of hepatic lipofuscin (Extended Data Fig. 3c,d and Fig. 5c). It is noteworthy that although the duration of TZP treatment was the same for both regimens (10 weeks), TZP treatment initiated concurrently with the start of HFD nearly completely abrogated lipofuscin accumulation (Extended Data Fig. 3c). In contrast, delayed TZP treatment, which started after 7 weeks of HFD, only reduced lipofuscin levels by 36% (Fig. 5c). In addition, a deeper analysis of hepatic lipofuscin levels across the study revealed that delayed TZP treatment failed to reduce the lipofuscin load below the levels observed at the time the treatment was initiated (area of SBB: ∼ 3.9% for 7 weeks of HFD + 10 weeks of TZP, Fig. 5c versus ∼ 1.8% for 7 weeks of HFD, Fig. 1f). These potentially suggested that TZP prevented lipofuscin formation, instead of being able to remove lipofuscin once it formed in the lysosome. With a significant reduction of lipofuscin, hepatic levels of CXCL5 (Extended Data Fig. 3e and Fig. 5d), neutrophil recruitment (Extended Data Fig. 3f and Fig. 5e), and NET formation in the liver (Extended Data Fig. 3g and Fig. 5f) were all dampened in the HFD-fed mice. TZP, however, had no effect on levels of lipid accumulation (Extended Data Fig. 4a,b).

**Fig. 5.**
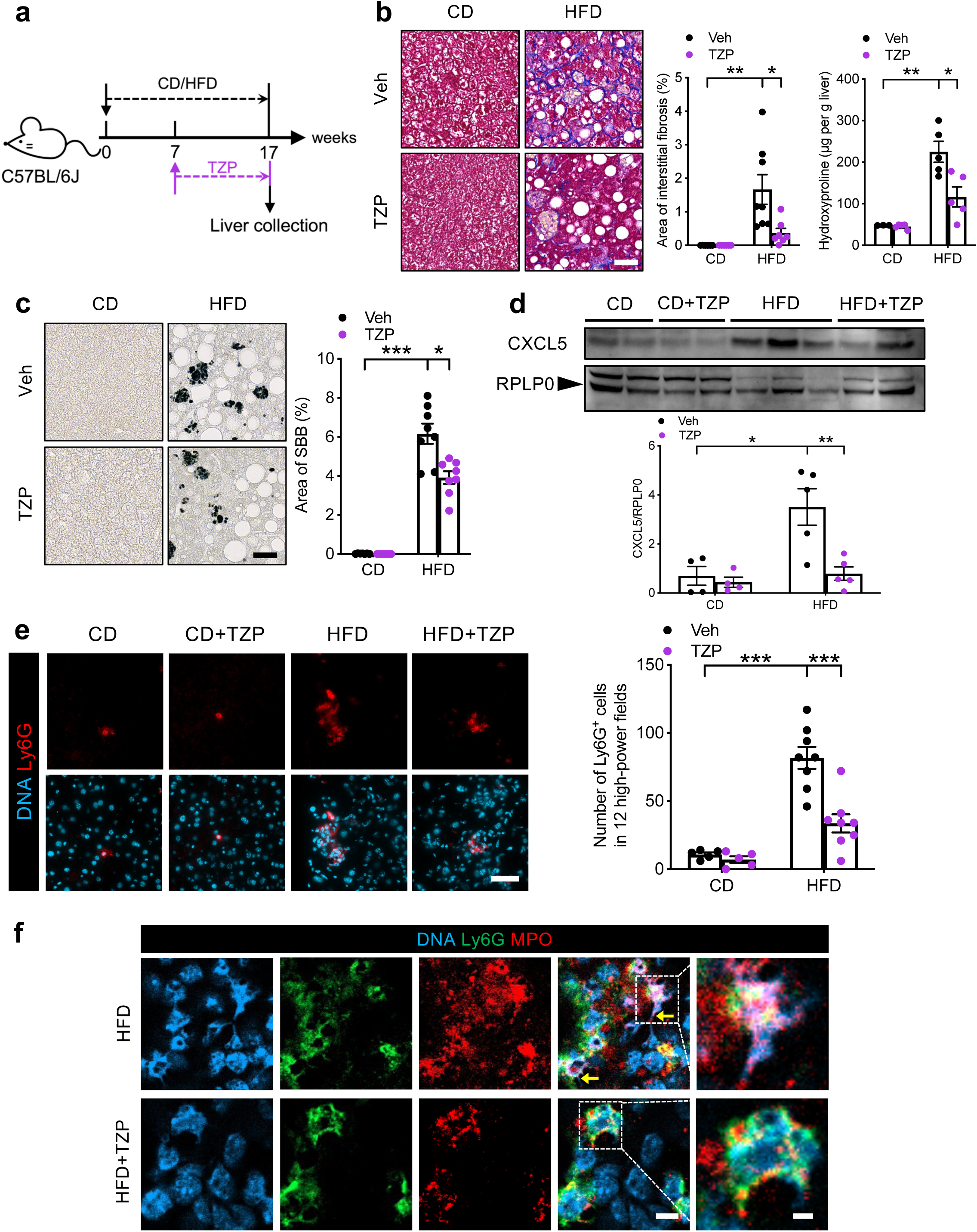
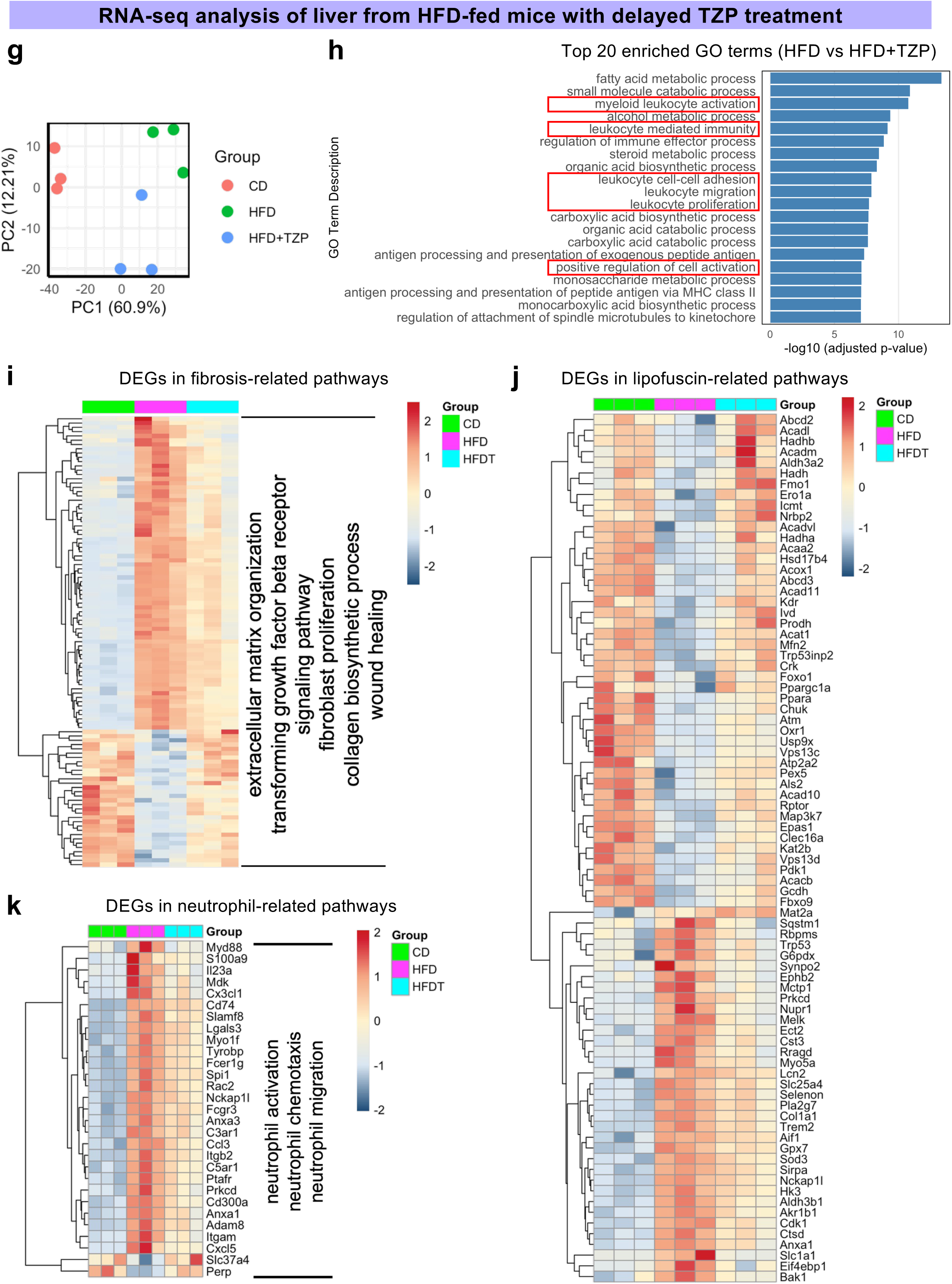
Tirzepatide (TZP) treatment at the mild liver fibrosis stage diminishes liver fibrosis severity by reducing hepatic lipofuscin load and the associated inflammation. **a-k,** Male C57BL/6J mice were fed CD or HFD for 7 weeks, followed by treatment with TZP or vehicle for the subsequent 10 weeks along with the respective diets (**a**), and bulk RNA-seq was performed on the liver tissue from these mice (*n* = 3 per group) (**g-k**). **b,** Representative images of liver interstitial fibrosis revealed by Masson’s trichrome (left) and quantification of area of interstitial fibrosis (%) (middle) on the FFPE liver sections. Level of liver fibrosis assessed by hydroxyproline content (µg per g liver) (right). For quantification of area of interstitial fibrosis (%), left to right on the bar chart, *n* = 8, 6, 8, 7. For hydroxyproline assay, left to right on the bar chart, *n* = 3, 5, 5, 5. Scale bar, 50 µm. **c,** Representative images of lipofuscin stained by SBB (left) and quantification of area of SBB (%) (right) of the liver sections. Left to right on the bar chart, *n* = 7, 7, 8, 8. Scale bar, 50 µm. **d,** Representative Western blots (top) and quantification of CXCL5 levels (bottom) in the liver. RPLP0 served as the loading control. Left to right on the bar chart, *n* = 4, 4, 5, 5. **e,** Immunofluorescence staining for neutrophil recruitment (Ly6G^+^ cells) in the liver (left) and quantification of neutrophils in 12 high-power fields (right). Left to right on the bar chart, *n* = 5, 5, 8, 8. Scale bar, 50 µm. Statistical comparisons were made using two-tailed unpaired t-test (**b,d,e**) or Mann-Whitney test (**c**). \**P* < 0.05, \*\**P* < 0.01 and \*\*\**P* < 0.001. Data are mean ± s.e.m. **f,** Single-plane confocal images of neutrophils and NETs in the liver of HFD-fed mice treated with vehicle or TZP. Top, yellow arrows indicate NETs; bottom, non-NETing neutrophils. Regions outlined by the white dotted boxes are magnified on the immediate right. Scale bars, 5 µm (left) and 2 µm (right). **g,** Principal component analysis of the bulk RNA-seq data from the liver of mice with different treatments. **h,** Top 20 enriched GO terms based on adjusted p-value in the liver of mice fed HFD with and without TZP treatment. **i-k,** Heatmap of DEGs in fibrosis-related pathways (extracellular matrix organization, transforming growth factor beta receptor signaling pathway, fibroblast proliferation, collagen biosynthetic process, wound healing) (**i**), lipofuscin-related pathways (autophagosome-lysosome fusion, response to oxidative stress, cellular response to oxidative stress, intrinsic apoptotic signaling pathway in response to oxidative stress, lipid oxidation, macroautophagy, TOR signaling) (**j**), and neutrophil-related pathways (neutrophil activation, neutrophil chemotaxis, neutrophil migration) (**k**).

### TZP reverses transcriptomic signatures related to fibrosis, neutrophil activation and lipofuscin accumulation in the liver of HFD-fed mice

Liver tissue of mice treated under both regimens of TZP were subjected to bulk RNA-seq analysis. Principal-component analysis revealed the distinctive pattern of transcriptome between HFD-fed mice with and without TZP treatments (Extended Data Fig. 5a and Fig. 5g). Pathways associated with fibrosis, including extracellular matrix structure and wound healing, as well as those related to immune regulation, such as leukocyte migration and myeloid leukocyte activation, were amongst the most enriched GO terms in the liver of HFD-fed mice compared to their counterparts with TZP treatment (Extended Data Fig. 5b and Fig. 5h). A comprehensive analysis of DEGs in key biological processes related to fibrosis demonstrated that gene expression in the liver of HFD-fed mice was largely reversed by TZP treatments to more CD-like (Extended Data Fig. 5c and Fig. 5i). Particularly, transcriptional levels of genes that encode extracellular matrix components (e.g., *Col1a1*, *Col1a2*, *Col3a1*) and enzymes regulating tissue remodeling (e.g., *Mmp12*, *Mmp2*, *Adamtsl2*) were upregulated in the liver by HFD and suppressed by TZP (Supplementary Table 1,2). Besides, TZP treatments reduced RNA signatures in the fibrogenic signaling pathways, including growth factors that promote fibroblast proliferation and differentiation (e.g., *Pdgfb*, *Tgfb1/2/3*), which were elevated by HFD (Supplementary Table 1,2). The overall inflammatory milieu was also modulated by TZP, with a notable reduction in the expression of genes involved in antigen presentation (e.g., *Cd74*) (Supplementary Table 1,2). Collectively, these findings suggested that TZP, in both preventive and therapeutic regimens, effectively normalizes a wide range of transcriptomic signatures related to fibrosis.

Since lipofuscin was associated with liver fibrosis, we also examined transcriptomic changes in lipofuscin-related pathways (Fig. 1e) after TZP treatments and found that the most relevant gene sets were related to autophagy, mitochondrial function, and oxidative stress (Extended Data Fig. 5d and Fig. 5j). Impaired autophagy, suggested by upregulation of *Trem2* and downregulation of *Foxo1*, was observed in the liver of HFD-fed mice (Extended Data Fig. 5d and Fig. 5j). After TZP treatments, expression patterns of these two genes were similar to those in CD-fed mice (Extended Data Fig. 5d and Fig. 5j), suggesting that enhanced autophagy might reduce lipofuscin accumulation^49^. Increased mRNA levels of genes related to hepatic fatty acid metabolism and mitochondrial biogenesis (e.g. *Acadvl*, *Hadha*, *Ppargc1a*) were observed in TZP-treated mice fed HFD (Extended Data Fig. 5d and Fig. 5j), implicating an improvement in mitochondrial function. Interestingly, HFD induced higher expression of antioxidant genes (e.g. *Sod3*, *Gpx7*, *Selenon*), which were reduced by both TZP treatment regimens (Extended Data Fig. 5d and Fig. 5j). This could be a compensatory response to counter the oxidative stress induced by HFD^33^; the exacerbated antioxidant defense was lowered when TZP normalized the oxidative microenvironment. Collectively, these results suggested that TZP restores the dysfunctional processes associated with lipofuscin accumulation in HFD.

TZP treatments also normalized DEGs related to pathways of neutrophil recruitment and activation (Extended Fig. 5e and Fig. 5k). Transcriptional levels of genes encoding chemokines and their receptors (e.g. *Cx3cl1, C5ar1*, *C3ar1*) that were potentially involved in neutrophil chemotaxis were downregulated by TZP, in addition to *Cxcl5* (Extended Fig. 5e and Fig. 5k). TZP treatments also suppressed expression of key players in neutrophil activation (e.g. *Lgals3*, *Fcgr3*, *Fcer1g*) (Extended Fig. 5e and Fig. 5k). Neutrophil adhesion was also dampened in the liver of TZP-treated mice, as reflected by a reduction in mRNA levels of *Itgam* and *Itgb2* (Extended Fig. 5e and Fig. 5k).

### TZP reverses cellular senescence induced by HFD

Cellular senescence or premature aging predisposes liver to chronic inflammation, which can result in liver fibrosis^50–52^. We sought to examine if HFD biologically aged the liver. Schaum et al. mapped molecular hallmarks of chronological aging across different organs^53^. By cross-referencing their findings with our dataset, aging-related Kyoto Encyclopedia of Genes and Genomes pathways were found to be significantly enriched in the liver of HFD-fed mice, and longevity regulating pathway appeared to be the most relevant in this HFD-induced liver fibrosis model (Extended Data Fig. 6a and Fig. 6a). While more aging-related DEGs were observed in mice fed HFD for 17 weeks (Fig. 6b) compared to those on HFD for 10 weeks (Extended Data Fig. 6b), DEGs associated with insulin/insulin-like growth factor 1 and mammalian target of rapamycin pathways were commonly altered (Extended Data Fig. 6b and Fig. 6b). Similar to those that decreased in chronological aging (e.g. *Igf1*, *Rptor*, *Foxo1*), HFD also downregulated their gene expression (Extended Data Fig. 6b and Fig. 6b). For those that increased with age, such as *Cryab*, HFD boosted their transcriptional levels as well (Extended Data Fig. 6b and Fig. 6b). Both regimens of TZP treatment were able to normalize part of the aging-related transcriptome (Extended Data Fig. 6c and Fig. 6c), suggesting a rejuvenating effect of TZP. DEGs in the liver of HFD-fed mice also highly overlapped with SenMayo gene panel that identified transcriptomic signatures of senescent cells^54^, and TZP treatments resulted in a gene expression pattern more similar to that of the CD-fed mice (Extended Data Fig. 6d and Fig. 6d). Concurrent treatment of TZP significantly reduced transcript levels of multiple genes encoding senescence-associated secretory phenotype (SASP) (e.g., *Mmp12*, *Mmp2* and *Spp1*) (Extended Data Fig. 6d), while delayed treatment only decreased *Mmp12* expression, showing a more limited effect (Fig. 6d).

**Fig. 6.**
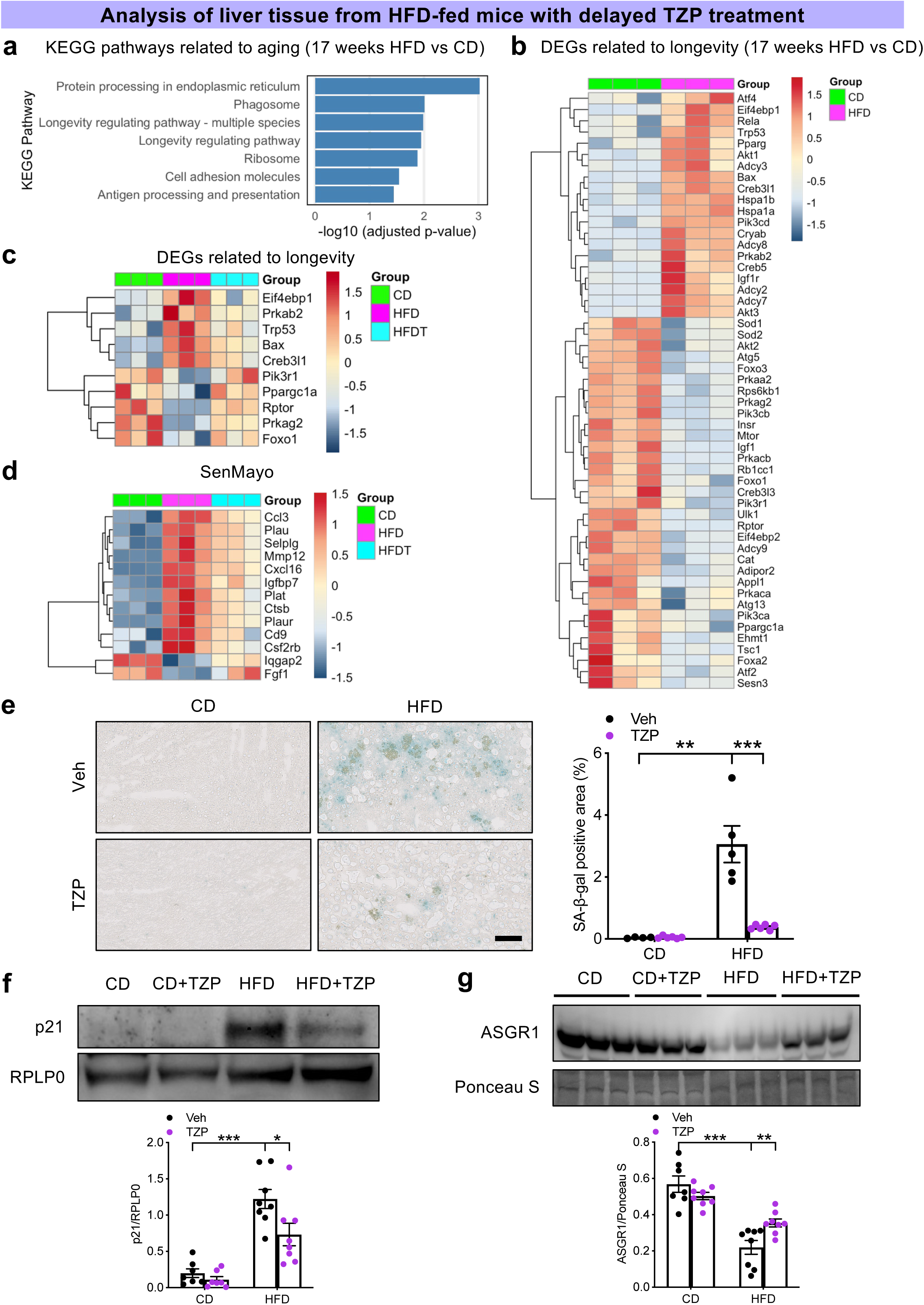
Delayed treatment of TZP reverses cellular senescence induced by HFD. Male C57BL/6J mice were fed CD or HFD for 7 weeks, followed by treatment of TZP or vehicle for the subsequent 10 weeks along with the respective diets (**a-g**) and bulk RNA-seq was performed on the liver tissue from these mice (*n* = 3 per group) (**a-d**). **a,** Aging-related Kyoto Encyclopedia of Genes and Genomes (KEGG) pathways enriched in the liver of mice fed HFD for 17 weeks compared to their CD-fed counterparts. **b,c,** Heatmap of DEGs in KEGG pathways related to longevity (longevity regulating pathway - multiple species, longevity regulating pathway) in the liver of mice without (**b**) and with (**c**) TZP treatment. **d,** Heatmap of DEGs in the liver transcriptome that overlaps with the SenMayo gene panel. **e,** Representative images of senescence-associated beta-galactosidase (SA-β-gal) staining (left) and quantification of SA-β-gal positive area (%) (right) on the liver sections. Left to right on the bar chart, *n* = 4, 6, 5, 6. Scale bar, 100 µm. **f,g,** Representative Western blots and quantification of levels of p21 (**f**) and ASGR1 (**g**). RPLP0 and Ponceau S-stained protein served as the loading controls. **f**, Left to right on the bar chart, *n* = 6, 6, 7, 7. **g,** Left to right on the bar chart, *n* = 7, 8, 8, 8 per group. Statistical comparisons were made using two-tailed unpaired t-test. \**P* < 0.05, \*\**P* < 0.01 and \*\*\**P* < 0.001. Data are mean ± s.e.m.

To confirm that TZP could reduce HFD-induced liver senescence, we examined multiple markers of cellular senescence with staining and Western blotting. One of the gold standards to identify senescent cells is to stain for senescence-associated beta-galactosidase (SA-β-gal)^55^. We observed that the liver of HFD-fed mice was positively stained with SA-β-gal (Extended Data Fig. 6e and Fig. 6e). Of note, the SA-β-gal signals were mostly observed in lipofuscin-heavy areas, which appeared yellowish-brown in the liver sections (Extended Data Fig. 6e and Fig. 6e). Intriguingly, cells that did not contain lipofuscin but were close to the lipofuscin-laden cells were also stained positive with SA-β-gal (Extended Data Fig. 6e and Fig. 6e), suggesting that lipofuscin-laden senescent cells could promote aging in neighboring cells through SASP, in line with the HFD-induced upregulation of SASP genes unveiled by the transcriptomic analysis. TZP treatments significantly reduced SA-β-gal staining in the liver of HFD-fed mice (Extended Data Fig. 6e and Fig. 6e). p21, a driver of cellular senescence by arresting the cell cycle^55^, was significantly upregulated in the liver of HFD-fed mice and was markedly decreased by TZP treatments (Extended Data Fig. 6f and Fig. 6f).

In addition, loss of cellular identity seen in aged cells^56^ was evident in hepatocytes of HFD-fed mice, and was effectively prevented by TZP treatments (Extended Data Fig. 6g and Fig. 6g). Hepatocyte polyploidization suggests terminal differentiation and cellular senescence^57^. We observed that majority of the lipofuscin-laden hepatocytes underwent polyploidization, displaying an average of 6 nuclei per cell, while hepatocytes without lipofuscin deposition typically contained 2 nuclei per cell (Extended Data Fig. 6h-j). Congruent with the reduction in lipofuscin-laden cells, number of hepatocytes undergoing polyploidization was reduced by 78% following TZP treatments (Extended Data Fig. 6k).

As senescent cells could escape immune surveillance and progress to cancer^58^, expression of markers relevant to liver cancer^59^ and stemness^60^ was also examined. Bulk RNA-seq analysis showed that carbamoyl phosphate synthetase 1 (encoded by *Cps1*, tumor suppressive) was decreased by HFD, while CD44 (encoded by *Cd44*, highly expressed in cancer stem cells) was elevated in the liver of HFD-fed mice (Extended Data Fig. 7a,b). Changes in these genes were reversed by TZP treatments (Extended Data Fig. 7a,b), suggesting that TZP might reduce cancer risk.

Similar to lipofuscin accumulation which was independent of NETosis, other HFD-induced senescence biomarkers, such as increased SA-β-gal activity (Extended Data Fig. 8a), elevated p21 expression (Extended Data Fig. 8b), and loss of hepatocyte identity (Extended Data Fig. 8c) were not rescued in the PAD4-deficient mice. These findings indicated that deficiency in NETosis did not prevent cellular senescence.

## Discussion

In this study, we unveiled a previously unrecognized inflammatory axis wherein lipofuscin-laden hepatocytes predispose neutrophils to NETosis that consequentially contributes to liver fibrosis. Intercepting NET formation directly or preventing lipofuscin accumulation by TZP both effectively reduce liver fibrosis. Surprisingly, TZP treatments also reverse hepatic senescence induced by HFD (Fig. 7). Our novel findings highlight the potential for TZP to combat liver fibrosis that is presented with lipofuscin.

**Fig. 7.**
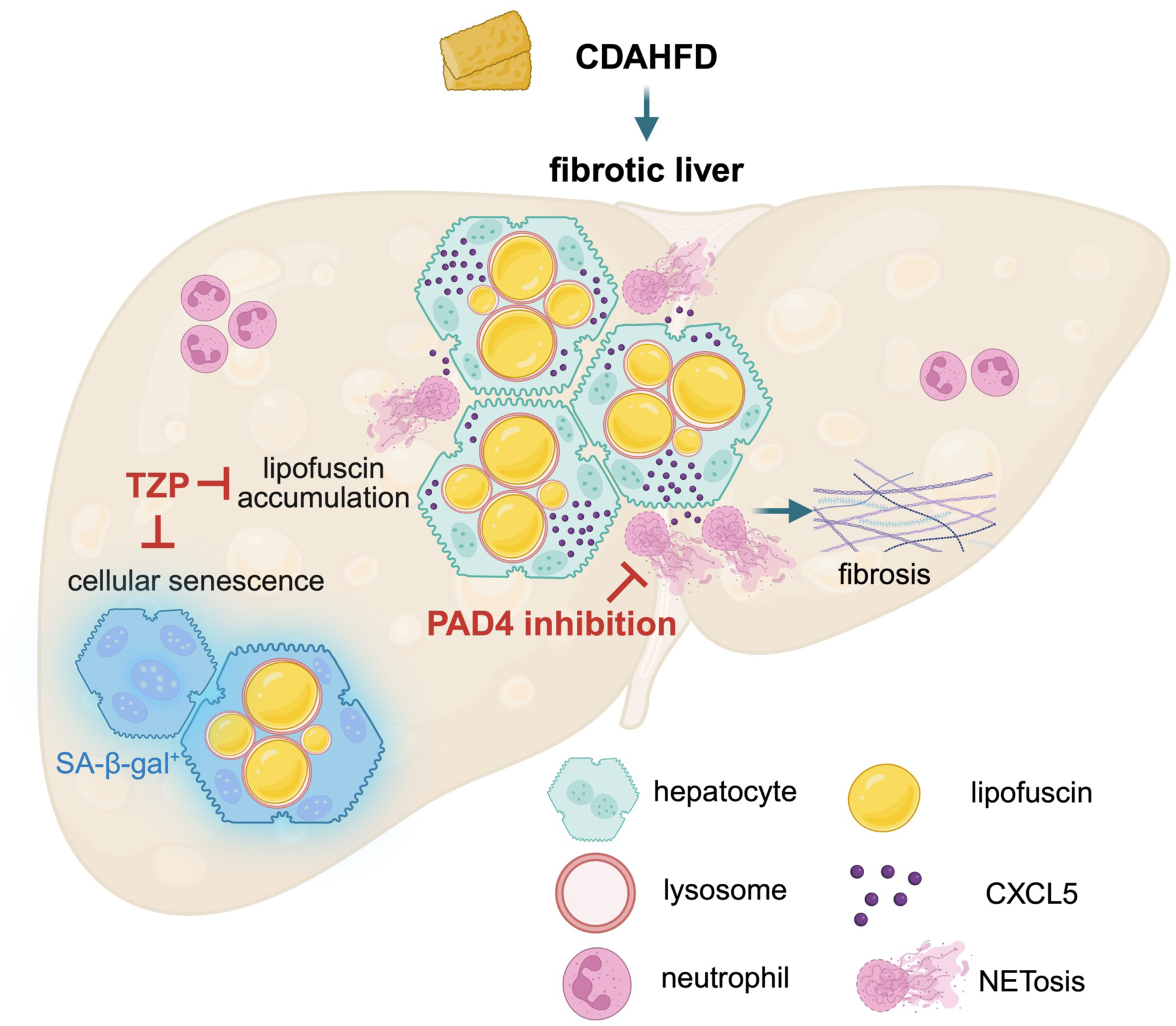
Schematic summary of how lipofuscin accumulation leads to liver fibrosis and the impact of TZP treatment. Choline-deficient L-amino acid-defined high-fat diet (CDAHFD) induces lipofuscin deposition in the lysosomes of hepatocytes. Lipofuscin-containing cells produce higher levels of CXCL5, which in turn enhances NET formation, eventually leading to liver fibrosis. Intercepting NETosis through direct inhibition of peptidylarginine deiminase 4 (PAD4) or through decreasing hepatic lipofuscin load by TZP treatment both effectively reduce liver fibrosis. TZP treatment provides additional therapeutic benefits by lessening hepatic cellular senescence. The illustration was created with BioRender.com.

While lipofuscin was conventionally regarded as a by-product of incomplete cellular catabolism without any pathological function^61^, several recent studies have revealed the deleterious effect of lipofuscin in different cell types^15,16,62,63^. Lipofuscin accumulating in the retinal pigment epithelium induces lysosome membrane permeabilization, which leads to necroptosis and retinal degeneration^15^. In aging cardiomyocytes, lipofuscin disrupts late-stage autophagy, resulting in impaired contractility^64^. Herein, we defined the critical pathogenic role of lipofuscin in liver disease. We mapped the relative timeframe of lipofuscin emergence and liver fibrosis, and found that lipofuscin accumulation preceded collagen deposition in the liver. The severity of liver fibrosis strongly and positively correlated with the hepatic lipofuscin load. Pathogenically, lipofuscin-laden hepatocytes produced CXCL5, which promoted neutrophil recruitment and NET release, resulting in liver fibrosis. Our findings revealed the upstream lipofuscin-NETosis axis, extending Xia et al.’s study which reports that NETs can induce cyclooxygenase-2-dependent prostaglandin E2 production in hepatic stellate cells that in turn generate extracellular matrix to fuel liver fibrosis in MASH models^65^. In our study, we showed that even lipofuscin accumulation and steatosis remained unaffected by NETosis inhibition as in the PAD4-deficient mice, liver fibrosis severity was remarkably reduced, suggesting a possible therapeutic approach that targets immune dysfunction in CLD.

Nonetheless, such promising pre-clinical findings cannot be easily translated as there are currently no PAD4 inhibitors that are clinically available. An alternative approach is the clearance of NETs using DNase 1 (dornase alfa). However, except for the DNA backbone of NETs, other cytotoxic components including histones and neutrophil elastase could not be effectively cleared by DNase 1^66^. To this end, our exciting findings of TZP shed light on new treatment possibilities. As the latest GIPR and GLP1R dual agonist approved for the treatment of type 2 diabetes^28^, TZP was shown to have additional benefits in MASH patients with liver fibrosis^29^. The combined use of individual GIPR and GLP1R agonists also reduces liver steatosis in a diabetic dyslipidemia mouse model^67^. However, these studies primarily focused on metabolic dysfunction-associated CLD and TZP’s effect on metabolism^68,69^. We showed that TZP decreased liver fibrosis by reducing the amount of hepatic lipofuscin and the associated NET formation, suggesting that TZP could be instrumental for CLD that presents with lipofuscin, beyond its application in MASH.

Two TZP treatment regimens (concurrent with the start of diets or delayed) were performed. While the original aim was to examine the effect of TZP on liver fibrosis early on (preventive approach) or delayed until mild fibrosis has taken place (therapeutic approach), the data interestingly provided clues on how TZP impacts on lipofuscin accumulation. Both concurrent and delayed TZP treatment regimens significantly lowered the hepatic lipofuscin load; however, the delayed treatment was not as effective as the concurrent treatment of the same duration in abolishing lipofuscin accumulation, nor could it reduce lipofuscin load to levels lower than those before the treatment. These data may suggest that TZP prevents the formation of lipofuscin, instead of facilitating its degradation. In fact, current perspectives on lipofuscin accumulation remain equivocal and impaired autophagy is one of the main proposed mechanisms^70^. Our bulk RNA-seq analysis uncovered compromised autophagy-related processes under HFD, which were restored by TZP treatments. This new data supports previous observations that rapamycin, an autophagy inducer, reduced lipofuscin deposition in the myocardium^49,64^. Given its profound effects on inhibiting lipofuscin accumulation, TZP could be a valuable experimental tool in understanding the complex regulation mechanisms of lipofuscin.

In addition to lipofuscin, TZP also effectively normalized other hepatic senescence signatures induced by HFD. Several studies support that aging and senescence promote the progression of liver disease^52,71^. Hepatic levels of p21 significantly correlate with fibrosis stage in human subjects^71^. The causal link between hepatocyte senescence and liver damage is demonstrated by a mouse study that removal of senescent cells using senolytic drugs reduces age-related liver steatosis^52^. However, there are also studies showing that senescence in hepatic stellate cells^72^ and liver sinusoidal endothelial cells^73^ prevents liver fibrosis. Hence, how senescence contributes to liver dysfunction appears highly dependent on the cell types that are aging. In the current study, majority of the lipofuscin-containing cells were hepatocytes (ASGR1^+^), supporting previous findings which showed that hepatocyte senescence facilitated liver pathogenesis^52,71^. According to these observations, senolytics may appear as an attractive treatment option for CLD. Nonetheless, their effect on hepatocyte senescence is not consistent. Dasatinib and quercetin, the widely recognized senolytics, not only fail to lessen hepatocyte senescence in drug-induced liver injury^74^, but also enhance obesity- and age-dependent liver disease^75^. TZP could thus be a novel and superior option to combat liver disease with lipofuscin accumulation and/or hepatic senescence. Our bulk RNA-seq analysis also unexpectedly revealed that TZP reversed several cancer-associated mRNA signatures present in the liver of HFD-fed mice. The finding is in line with Sagy et al.’s study which reports that obese and diabetic patients treated with GLP1R agonists are associated with a lower cancer incidence when compared to individuals who received bariatric metabolic surgery, even the latter has a stronger effect in decreasing body weight^76^. In fact, our findings may suggest novel causation insights to explain Sagy et al.’s observation – Since senescent hepatocytes may evade immune surveillance and become carcinogenic^58^, TZP or other GLP1R agonists that can reduce cellular senescence and inflammation will indirectly lower cancer risk. Of note, we found that amount of food intake was unaffected in TZP-treated mice (Extended Data Fig. 9), possibly attributed to the low injection frequency compared to other studies^77,78^. This agrees with clinical studies in which only a small percentage of patients (6-16%) experience decreased appetite with TZP treatment^28,29,79^. While more investigations regarding long-term TZP treatment are underway, it appears promising that TZP could be a new therapeutic option for senescence-related diseases.

One main challenge that we encountered in the study was the strong autofluorescence of lipofuscin that limited the investigation. While the autofluorescence can be blocked using SBB on liver sections to enable immunofluorescence staining and microscopy, such quenching protocol is not applicable for flow cytometry. Therefore, defining subtypes of neutrophils (e.g., NETting versus non-NETing neutrophils in the liver, blood neutrophils versus neutrophils recruited to the liver) and their characteristics in the homogenates could not be achieved by conventional flow cytometry as the strong autofluorescent signals interfered all fluorescence channels. To study neutrophil heterogeneity in-depth, full-spectrum flow cytometry is essential to capture the real signals (analysis) and to isolate them for functional characterization (sorting) by unmasking the autofluorescence background.

In conclusion, our study has defined the pathogenic role of lipofuscin in liver fibrosis. Lipofuscin-containing cells induce NET formation; inhibition of which effectively reduces liver fibrosis. TZP treatment targets lipofuscin accumulation and lessens the associated neutrophil activation. As it also reverses a battery of cellular senescence signatures, TZP could become a new generation of senotherapeutics for liver disease and beyond.

## Methods

### Cell culture

Human promyelocytic leukemia cell line HL-60 (ATCC CCL-240) was cultured in Roswell Park Memorial Institute (RPMI) 1640 medium (Gibco, 11875085) supplemented with 15% heat-inactivated fetal bovine serum, 1% penicillin/streptomycin and 25 mM HEPES (thereafter referred to as complete RPMI) at 37°C and 5% CO_2_. For differentiation towards neutrophil-like cells (dHL-60), 1.3% dimethyl sulfoxide (DMSO, Sigma-Aldrich, D2653) was added to the complete RPMI. The cell line was routinely tested to be free from mycoplasma.

### Animals

Experimental protocols of the study were reviewed and approved by Nanyang Technological University Institutional Animal Care and Use Committee (Protocol A19090 and A24088). C57BL/6J mice were purchased from InVivos (Singapore) and *Vav1-Cre Padi4^fl/fl^* were bred in-house. Only male mice were included in this study and all mice were used at 8 to 10 weeks of age. Mice were housed under specific pathogen-free conditions at 25°C with free access to diet and water.

To investigate fibrosis development and lipofuscin emergence in the liver, C57BL/6J mice were fed choline-deficient L-amino acid-defined high-fat diet (CDAHFD, later abbreviated as HFD, Research Diets Inc., A06071309i) or an ingredient-matched control diet (L-amino acid diet with normal levels of methionine and choline, later abbreviated as CD, Research Diets Inc., A06071322i) for 2, 7 and 10 weeks.

To investigate the role of NETs in liver fibrosis, mice defective in NETosis (*Vav1-Cre Padi4^fl/fl^*) and their control (*Padi4^fl/fl^*) were fed CD or HFD for 10 weeks.

To examine whether TZP could prevent the onset of liver fibrosis development, C57BL/6J mice were fed CD or HFD with concurrent treatment of TZP (Selleck Chemicals, P1206, subcutaneous injection, 30 nmol/kg) or vehicle (20 mM Tris-HCl, 0.05% Tween-80, pH 8.0) every 3 days for 10 weeks. To investigate whether delayed treatment of TZP could also alleviate liver fibrosis, C57BL/6J mice were fed CD or HFD for 7 weeks, followed by TZP treatment (30 nmol/kg, subcutaneous) or vehicle every 3 days for an additional 10 weeks along with the respective diets.

### Mouse liver collection and processing

After anesthesia, liver was perfused via the inferior vena cava using 20 mL of perfusion buffer (10 mM EDTA and 5 mM HEPES in 1xPBS) followed by 10 mL of 1xPBS. A small incision was made in the portal vein during perfusion to let out the perfusate. For histological analysis, autofluorescence imaging, senescence-associated beta-galactosidase (SA-β-gal) staining and immunofluorescence staining, liver was processed into formalin-fixed paraffin-embedded (FFPE) sample, or embedded in optimal cutting temperature (OCT) compound. For Western blotting and hydroxyproline assay, liver was snap-frozen in liquid nitrogen and kept at −80°C. Liver samples for bulk RNA sequencing were stored in RNAlater (Sigma-Aldrich, R0901) at −80°C until further procedures.

### Histology

FFPE liver samples were cut into 5 μm sections. The sections were then deparaffinized for hematoxylin and eosin staining, Masson’s trichrome staining, and Sudan Black B (SBB) staining. Trichrome Stain Kit (Sigma-Aldrich, HT15) was used for Masson’s trichrome staining per the manufacturer’s instruction. For SBB staining, deparaffinized liver sections were put into 0.7% SBB solution (Sigma-Aldrich, 199664) for 8 mins, followed by rinses in 70% ethanol, 50% ethanol and water before mounting. OCT-embedded liver samples were cut into 10 μm sections for Oil Red O (ORO) staining as previously described^80^. Images were acquired on a slide scanner (Carl Zeiss, Axio Scan.Z1) using the Plan Apochromat 20x/0.8 lens. Area of interstitial fibrosis (%), area of ORO (%) and area of SBB (%) were quantified from 20 to 40 high-power fields using ImageJ (v2.14.0).

### Autofluorescence imaging for lipofuscin

FFPE liver sections were used for autofluorescence imaging for lipofuscin. Images were acquired on a slide scanner (Carl Zeiss, Axio Scan.Z1) using a Plan-Apochromat 40x/0.95 Korr M27 objective. An LED-Module 385 nm excitation source was applied with 100% intensity and 20.01 ms exposure time. The excitation and emission wavelengths were 353 nm and 465 nm, respectively.

### SA-β-gal staining

Cryopreserved liver sections were fixed in 0.5% glutaraldehyde and washed in 1x PBS containing 1 mM MgCl_2_ (pH 5.5). Thereafter, freshly prepared X-gal solution (Gold Biotechnology, X4281C) was applied to sections for 6 hours at 37°C. Post-fixation was conducted using 4% paraformaldehyde. Images were acquired on a slide scanner (Carl Zeiss, Axio Scan.Z1) using the Plan Apochromat 20x/0.8 lens. Color deconvolution using Alcian blue & H vector was applied to images, and positive area of SA-β-gal staining (%) was quantified from 20 high-power fields through thresholding analysis using ImageJ (v2.14.0).

### Immunofluorescence staining

Sections from FFPE liver samples were deparaffinized and subjected to heat-induced antigen retrieval using Tris-EDTA buffer (pH 9.0) for staining ASGR1. Immunofluorescence staining of other proteins was performed on OCT-embedded liver sections after fixation in zinc fixative overnight. Sections were then put into 0.1% SBB solution for 30 minutes to block autofluorescence from lipofuscin. After permeabilization and blocking of non-specific binding, sections were stained with primary antibodies (rabbit anti-mouse ASGR1, Proteintech, 11739-1-AP, 1:250; rabbit anti-mouse CXCL5, Bioss, bs-2549, 1:200; rabbit anti-human myeloperoxidase, Dako, A0398, 1:500; rat anti-mouse Ly-6G, BD Bioscience, 551459, 1:500; rat anti-mouse LAMP1, Abcam, ab25245, 1:200) overnight at 4°C and appropriate secondary antibodies thereafter.

Z-stack images were obtained for slides stained with LAMP1 or ASGR1 using a laser scanning confocal microscope (Carl Zeiss, LSM 800) with the Plan Apochromat 63x/1.4 oil lens. Maximum intensity projection images were compiled from z-stack optical sections for LAMP1 and ASGR1 staining. Single-plane images were acquired for slides stained for myeloperoxidase and Ly6G to detect neutrophil extracellular traps (NETs) or ASGR1 using a laser scanning confocal microscope (Carl Zeiss, LSM 800) with the Plan Apochromat 63x/1.4 oil lens. Slides stained for Ly6G and CXCL5 were imaged with an inverted fluorescence microscope (Carl Zeiss, Axio Observer 7) using the Plan Apochromat 63x/1.4 oil lens. Number of Ly6G-positive cells in the liver sections was quantified from 12 high-power fields to determine neutrophil recruitment into the liver.

### Western blotting

Snap-frozen liver samples were homogenized in mammalian protein extraction reagent (Thermo Scientific, 78501) containing protease and phosphatase inhibitor (Thermo Scientific, 78441). Protein quantification was performed using Pierce™ BCA protein assay kit (Thermo Scientific, 23225). Eighty micrograms of protein were loaded onto 4-12% Bolt™ Bis-Tris Plus gel (Invitrogen, NW04120BOX) or 12% Bolt™ Bis-Tris Plus gel (Invitrogen, NW00122BOX) and transferred to PVDF membrane (Invitrogen, IB24002) using iBlot™ 2 Dry Blotting System (Invitrogen, IB21001). Membrane was blocked in 5% non-fat milk, and incubated with primary antibodies (rabbit anti-mouse ASGR1, Proteintech, 11739-1-AP, 1:1000; rabbit anti-mouse CXCL5, Bioss, bs-2549, 1:1000; rabbit anti-mouse RPLP0, Proteintech, 11290-2-AP, 1:1000; rabbit anti-mouse p21, Abcam, ab188224, 1:1000) at 4°C overnight and subsequently with goat anti-rabbit IgG (H+L) horseradish peroxidase (HRP)-conjugated antibody (Bio-Rad Laboratories, 1706515) for 2 hours at room temperature. The blots were developed using Sirius HRP substrate (Advansta, K-12043-D20). The band intensity was quantified using ImageJ (v2.14.0). RPLP0 or Ponceau S staining was used as a loading control.

### Hydroxyproline assay

Hepatic collagen content was measured using Hydroxyproline Assay Kit (Sigma-Aldrich, MAK008) according to manufacturer’s protocol. Briefly, snap-frozen liver samples were homogenized in water and hydrolyzed in 37% HCl to extract hydroxyproline. Subsequently, chloramine T was added to convert the hydroxyproline into pyrrole, which reacted with 4-(dimethylamino)benzaldehyde mixture to produce a chromophore. Absorbance was measured at 560 nm.

### Bulk RNA sequencing and analysis

Liver samples preserved in RNAlater were sequenced using the Illumina NovaSeq PE150 platform (Novogene, Singapore). Five hundred ng RNA per liver sample was collected to analyze the transcriptome. Raw data (raw reads) of fastq format were first processed through fastp software. Clean data (clean reads) were then obtained by removing reads containing adapter, reads containing poly-N and low-quality reads from raw data. Reads were aligned to mouse genome reference (GRCm39/mm39, Ensembl 107) using HISAT2 (v2.0.5)^81^. Read counts and Fragments Per Kilobase of transcript per Million mapped reads (fpkm) were quantified using featureCounts (v1.5.0-p3)^82^. Data quality control, mapping and quantification of gene expression level were provided by Novogene. Differentially expressed genes were identified using DESeq2 (v1.46.0)^83^. Gene Ontology (GO) enrichment analysis and Kyoto Encyclopaedia of Genes and Genomes (KEGG) analysis were performed using ClusterProfiler (v4.14.6)^84^. Heatmap was generated using *pheatmap* (https://github.com/raivokolde/pheatmap) and plots were generated using R *ggplot2* (v3.5.1)^85^. Sets of analysis can be found in Supplementary Table 3.

### Cell-based NETosis assay

To assess NETosis in mouse neutrophils, around 1 mL of blood was collected at the retro-orbital plexus after anesthesia. Blood neutrophils were isolated using discontinuous Percoll gradients (Cytiva, 17544502) and neutrophils were collected at 69%/78% interface^21^. Ten thousand cells/well were seeded in a 96-well plate. The neutrophils were stimulated with 10 µg/mL lipopolysaccharide (LPS) from *Klebsiella pneumoniae* (Sigma-Aldrich, L4268) or 4 µM ionomycin (Invitrogen, I24222) for 4 hours.

To examine the effect of CXCL5 on NETosis, HL-60 were differentiated with DMSO for 6 days, and then exposed to 50 ng/mL recombinant human CXCL5 (PeproTech, 300-22) for 24 hours. Ten thousand cells/well were seeded in a 96-well plate. Cells were stimulated with 4 µM ionomycin or 50 µM oleic acid for 4 hours in the presence of 50 ng/mL CXCL5.

After stimulation, mouse neutrophils or dHL-60 were fixed in 2% paraformaldehyde containing Hoechst 33342 overnight at 4°C. Twenty non-overlapping images for each condition were acquired on an inverted fluorescence microscope (Carl Zeiss, Axio Observer 7) using the LD-Plan Neofluar 20x/0.4 lens for quantification of NETs.

### Measurement of H4Cit^high^ cells

Differentiated HL-60 cells were exposed to CXCL5 as described above. Cells were stimulated with 4 µM ionomycin for 1 hour, and then fixed in 2% paraformaldehyde containing Hoechst 33342 overnight at 4°C. Cells were permeabilized, blocked, and stained with anti-histone H4 (citrulline 3) (Sigma-Aldrich, 07-596) and subsequently with Alexa Fluor 488 goat anti-rabbit IgG (H+L) cross-adsorbed secondary antibody (Invitrogen, A-11008). Twenty non-overlapping images for each condition were acquired on an inverted fluorescence microscope (Carl Zeiss, Axio Observer 7) using the LD-Plan Neofluar 20x/0.4 lens. Images were converted to an 8-bit format and appropriate thresholding based on cell size and fluorescent intensity was applied to determine H4Cit^high^ cells. Analysis was conducted using ImageJ (v2.14.0).

### Statistical analysis

Data are mean ± s.e.m. Statistical analysis was performed using GraphPad Prism version 9.0 (GraphPad Software). Data normality was tested using Shapiro-Wilk test or Kolmogorov-Smirnov test. Analysis of two groups was performed with two-tailed unpaired *t* test (if data is normally distributed) or Mann-Whitney test (if data is not normally distributed). P < 0.05 is considered statistically significant.

## Acknowledgements

The study was supported by Nanyang Assistant Professorship (The Lee Kong Chian School of Medicine, Nanyang Technological University Singapore Start-Up Grant), the Singapore Ministry of Health’s National Medical Research Council under its Open Fund - Individual Research Grant (MOH-001495), and Vascular Research Initiative (022415-00001, Lee Kong Chian School of Medicine Strategic Academic Initiative) (to S.L.W.). L.D.W. was supported by the Nanyang President’s Graduate Scholarship from Nanyang Technological University Singapore. F.C. is supported by Nanyang Technological University Research Scholarship.

## Author contributions

SL Wong conceptualized the research. F Chen, LD Wang, T Wuestefeld and SL Wong designed the experiments. F Chen, Q Nam and F Chammas conducted experiments, acquired and analyzed data. F Chen, Q Nam, LD Wang, F Chammas, T Wuestefeld and SL Wong interpreted data. BTK Lee provided critical guidance on RNA-seq data analysis and interpretation. SL Wong acquired funding for the research and supervised the study. F Chen and SL Wong wrote the manuscript. All authors critically reviewed the manuscript and approved the manuscript for submission.

## Data availability

Bulk RNA-seq data will be made available on NCBI GEO upon the acceptance of the manuscript.

---

**Extended Data Fig. 1:**
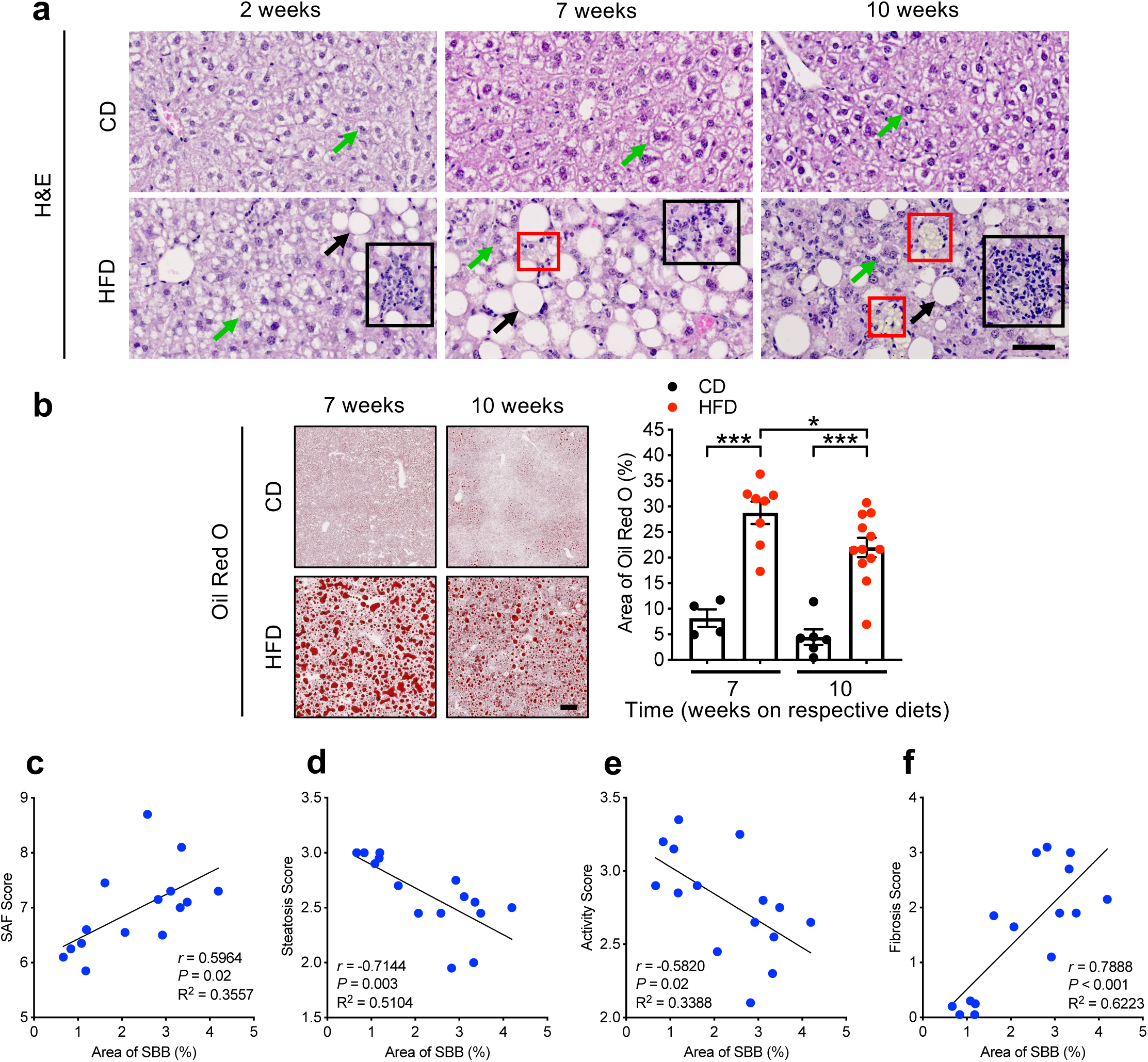
HFD induces other pathological features of chronic liver disease. **a,** Representative images of H&E-stained liver sections from mice fed the respective diets for up to 10 weeks, showing hepatocyte ballooning (green arrows), steatosis (black arrows), immune cell infiltration (black squares) and lipofuscin accumulation (red squares). Scale bar, 50 µm. **b,** Representative images of Oil Red O (ORO)-stained liver sections (left) and quantification of area of ORO (%) (right) of mice fed mice fed CD or HFD for 7 and 10 weeks. Left to right on the bar chart, *n* = 4, 8, 6, 12. Scale bar, 200 µm. Statistical comparison was made using two-tailed unpaired t-test. \**P* < 0.05 and \*\*\**P* < 0.001. Data are mean ± s.e.m. **c-f,** Pearson correlation analysis performed between area of SBB (%) versus steatosis, activity, and fibrosis (SAF) score (**c**), steatosis score (**d**), activity score (**e**), and fibrosis score (**f**). *n* = 5 for 7-week HFD group; *n* = 10 for 10-week HFD group.

**Extended Data Fig. 2:**
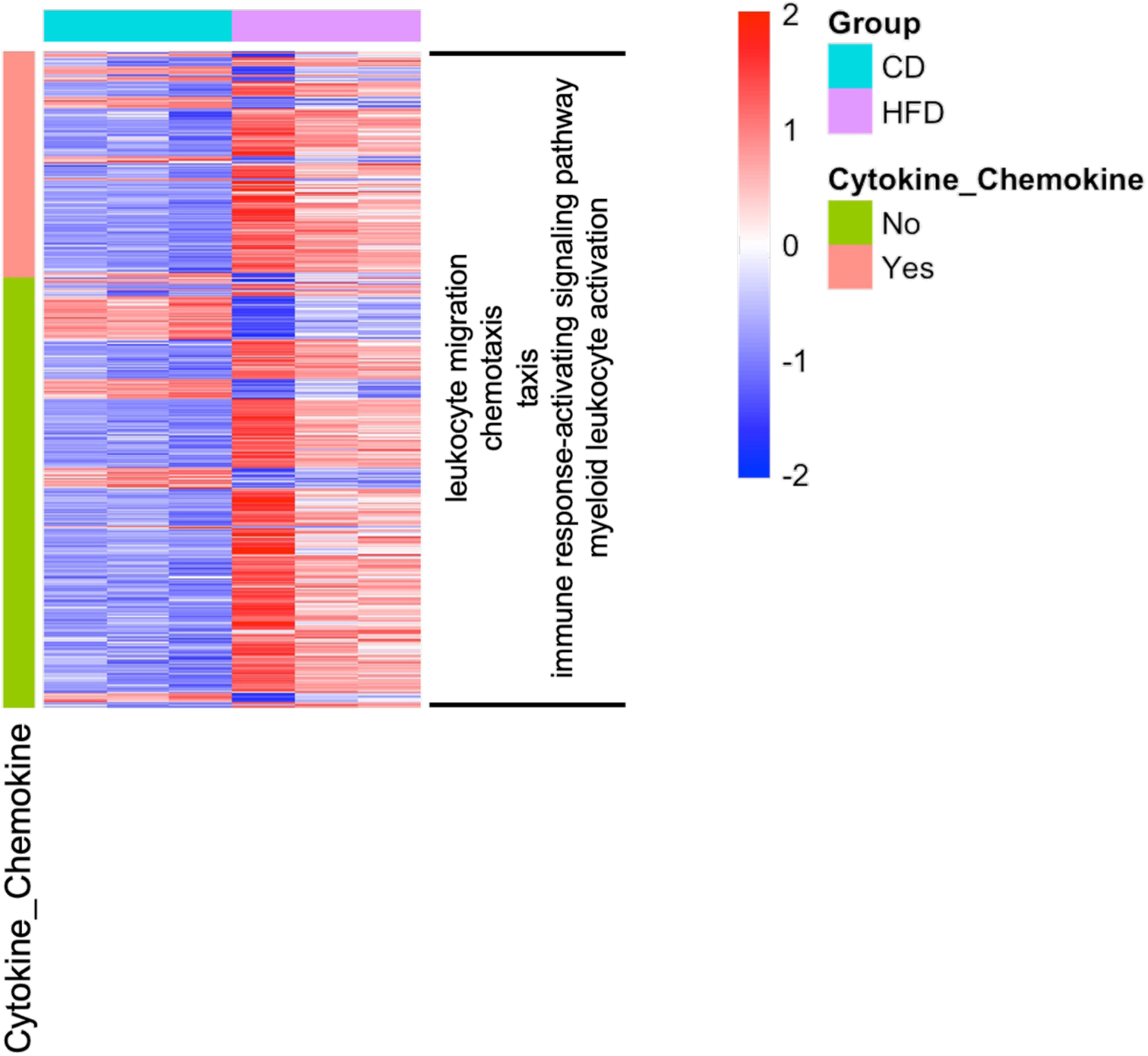
One-third of DEGs in top 20 enriched GO terms related to immune regulation are identified as cytokine/chemokine-related genes. Heatmap of DEGs in top 20 GO terms related to immune regulation (i.e., leukocyte migration, chemotaxis, taxis, immune response-activating signaling pathway, myeloid leukocyte activation) in the liver from mice fed HFD compared to mice fed CD for 10 weeks based on bulk RNA-seq.

**Extended Data Fig. 3:**
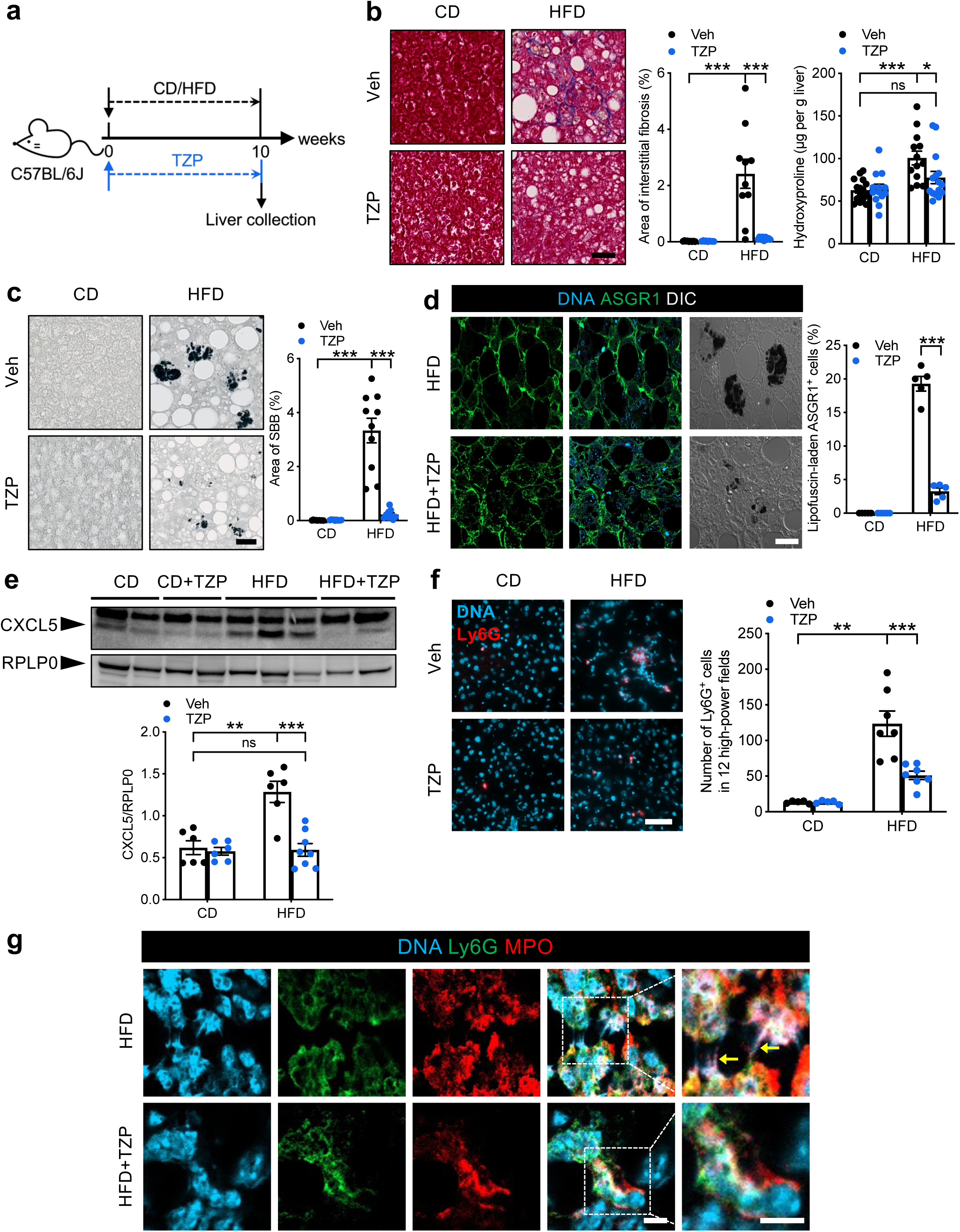
Concurrent treatment of TZP decreases liver fibrosis by reducing hepatic lipofuscin accumulation and the associated immune activation. **a-g,** Male C57BL/6J mice were fed CD or HFD with concurrent treatment of TZP or vehicle for 10 weeks (**a**). **b,** Representative images of liver interstitial fibrosis revealed by Masson’s trichrome (left) and quantification of area of interstitial fibrosis (%) (middle) on FFPE liver sections. Level of liver fibrosis assessed by hydroxyproline content (µg per g liver) (right). For quantification of area of interstitial fibrosis (%), left to right on the bar chart, *n* = 9, 9, 10, 9. For hydroxyproline assay, left to right on the bar chart, *n* = 15, 13, 13, 15. Scale bar, 50 µm. **c,** Representative images of lipofuscin stained by SBB (left) and quantification of area of SBB (%) (right). Left to right on the bar chart, *n* = 9, 10, 10, 9. Scale bar, 50 µm. **d,** Single-plane confocal images of hepatocytes (ASGR1^+^) (left) and quantification of lipofuscin-laden hepatocytes (%) (right) in the liver sections. *n* = 5 per group. Scale bar, 20 µm. **e,** Representative Western blots (top) and quantification (bottom) of CXCL5 levels in the liver. RPLP0 served as the loading control. Left to right on the bar chart, *n* = 6, 6, 6, 8. **f,** Immunofluorescence staining for neutrophil recruitment (Ly6G^+^ cells) in the liver (left) and quantification of neutrophils in 12 high-power fields (right). Left to right on the bar chart, *n* = 5, 5, 7, 7. Scale bar, 50 µm. Statistical comparisons were made using two-tailed unpaired t-test (**b-middle,c,d,e,f**) or Mann-Whitney test (**b-right**). \**P* < 0.05, \*\**P* < 0.01, \*\*\**P* < 0.001, ns, not significant. Data are mean ± s.e.m. **g,** Single-plane confocal images of neutrophils and NETs in the liver sections from HFD-fed mice treated with vehicle or TZP. Top, yellow arrows indicate NETs; bottom, non-NETing neutrophils. Regions outlined by white dotted boxes are magnified on the immediate right. Scale bars, 5 µm.

**Extended Data Fig. 4:**
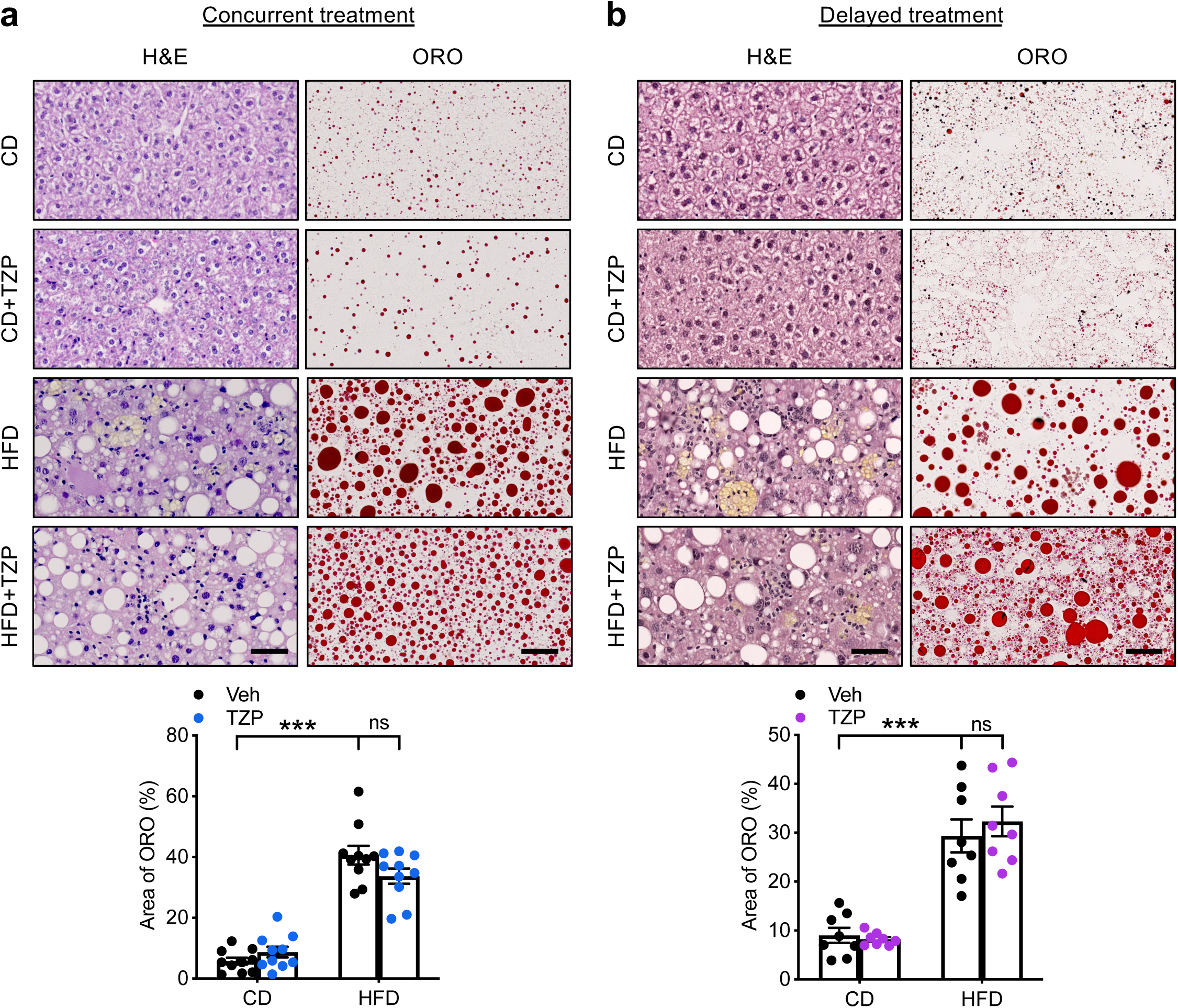
Treatment of TZP does not affect hepatic steatosis. **a,b** Representative images of H&E and ORO staining (top) and quantification of area of ORO (%) (bottom) of liver sections from mice with concurrent (**a**) or delayed (**b**) TZP treatment. *n* = 10 per group (**a**) and *n* = 8 per group (**b**). Scale bars, 100 µm (H&E images) and 50 µm (ORO images). Statistical comparisons were made using two-tailed unpaired t-test (**a,b**). \*\*\**P* < 0.001, ns, not significant. Data are mean ± s.e.m.

**Extended Data Fig. 5:**
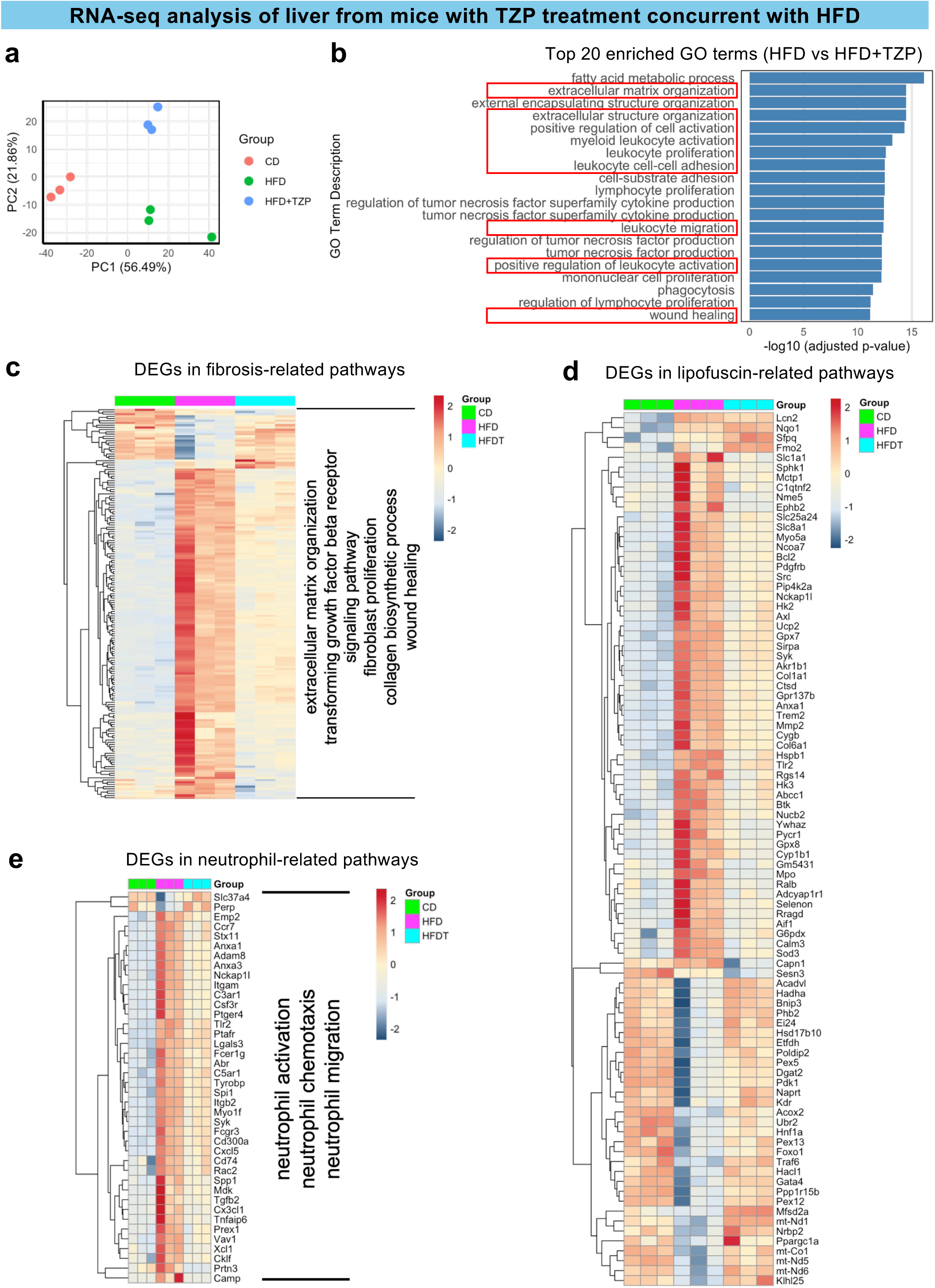
Concurrent treatment of TZP reverses transcriptomic signatures in fibrosis-, lipofuscin- and neutrophil-related pathways in HFD-fed mice. **a-e,** Male C57BL/6J mice were fed CD or HFD with concurrent treatment of TZP or vehicle for 10 weeks and bulk RNA-seq was performed on the liver from these mice (*n* = 3 per group). **a,** Principal component analysis of the bulk RNA-seq data from the liver of mice with different treatments. **b,** Top 20 enriched GO terms based on adjusted p-value in the liver of mice fed HFD with and without TZP treatment. **c-e,** Heatmap of DEGs in fibrosis-related pathways (extracellular matrix organization, transforming growth factor beta receptor signaling pathway, fibroblast proliferation, collagen biosynthetic process, wound healing) (**c**), lipofuscin-related pathways (response to oxidative stress, cellular response to oxidative stress, intrinsic apoptotic signaling pathway in response to oxidative stress, lipid oxidation, macroautophagy, TOR signaling) (**d**), and neutrophil-related pathways (neutrophil activation, neutrophil chemotaxis, neutrophil migration) (**e**).

**Extended Data Fig 6:**
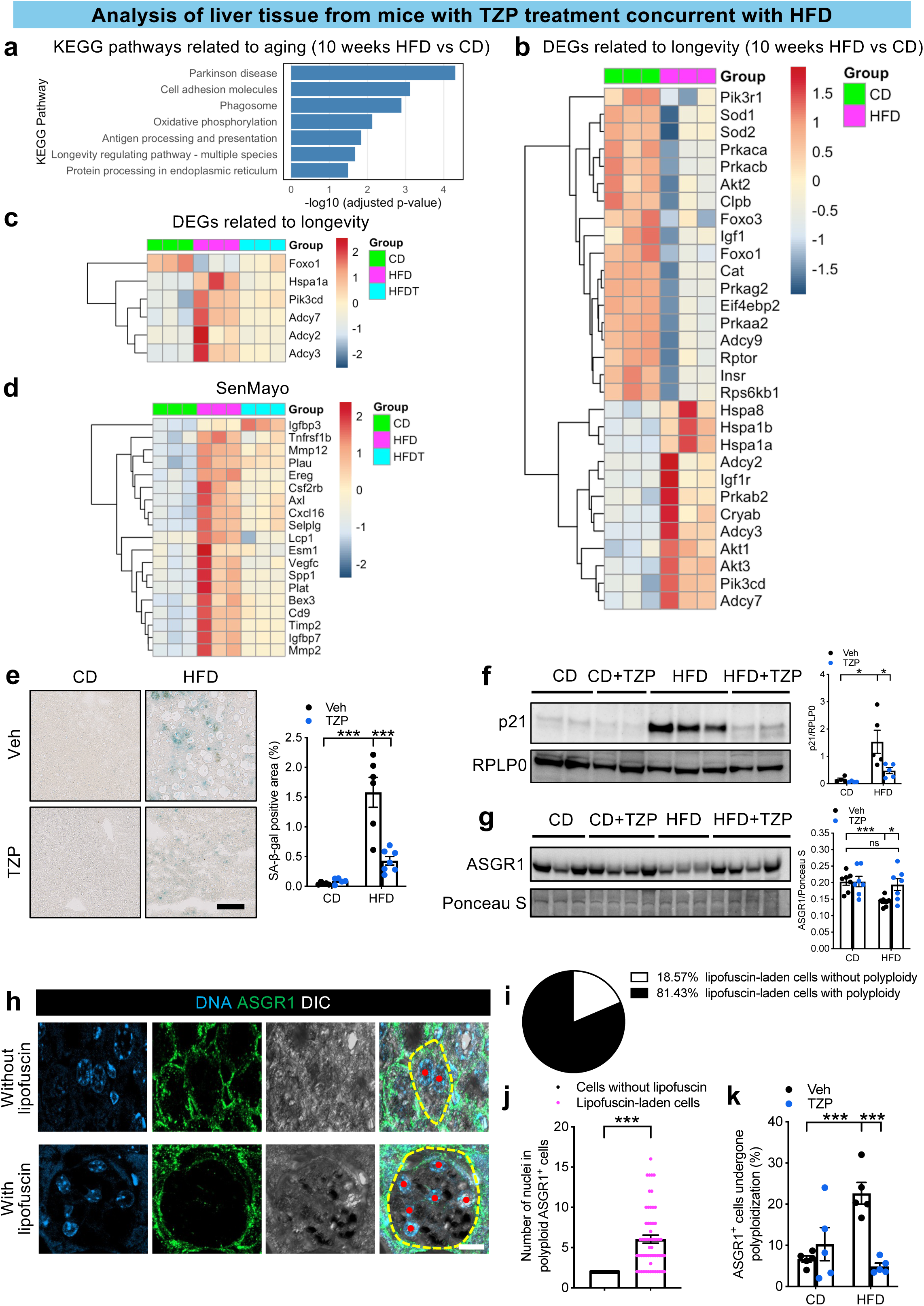
Concurrent treatment of TZP reverses cellular senescence induced by HFD. Male C57BL/6J mice were fed CD or HFD with concurrent treatment of TZP or vehicle for 10 weeks (**a-k**) and bulk RNA-seq was performed on the liver tissue from these mice (*n* = 3 per group) (**a-d**). **a,** Aging-related KEGG pathways enriched in the liver of mice fed HFD compared to mice fed CD for 10 weeks. **b,c,** Heatmap of DEGs in KEGG pathways related to longevity (longevity regulating pathway - multiple species) in the liver of mice without (**b**) and with (**c**) TZP treatment. **d,** Heatmap of DEGs in the liver transcriptome that overlaps with the SenMayo gene panel. **e,** Representative images of senescence-associated beta-galactosidase (SA-β-gal) staining (left) and quantification of SA-β-gal positive area (%) (right) on the liver sections. Left to right on the bar chart, *n* = 5, 5, 6, 7. Scale bar, 100 µm. **f,g,** Representative Western blots and quantification of the levels of p21 (**f**) and ASGR1 (**g**) in the liver. **f**, Left to right on the bar chart, *n* = 4, 4, 5, 5. **G**, *n* = 7 per group. RPLP0 (**f**) and Ponceau S-stained protein (**g**, used instead of RPLP0 since molecular size of RPLP0 was similar to the target band ASGR1) served as the loading controls. **h,** Representative images of hepatocytes (ASGR1^+^) without (top) and with (bottom) lipofuscin. Maximum intensity projection was applied to Z-stack images for slides stained with ASGR1. Cell membrane of hepatocyte is delineated by yellow dotted lines and nuclei are indicated by red dots. Scale bar, 10 µm. **i,** Pie chart showing percentage of lipofuscin-laden cells with (57 out of 70 cells, in black) or without polyploidy (13 out of 70 cells, in white). **j,** Number of nuclei in polyploid hepatocytes without (53 cells examined) or with (57 cells examined) lipofuscin. Each point represents one ASGR1^+^ cell. **k,** Quantification of ASGR1^+^ cells that underwent polyploidization (%). *n* = 5 per group. Statistical comparisons were made using two-tailed unpaired t-test (**e-g,k**) or Mann-Whitney test (**j**). \**P* < 0.05 and \*\*\**P* < 0.001, ns, not significant. Data are mean ± s.e.m.

**Extended Data Fig. 7:**
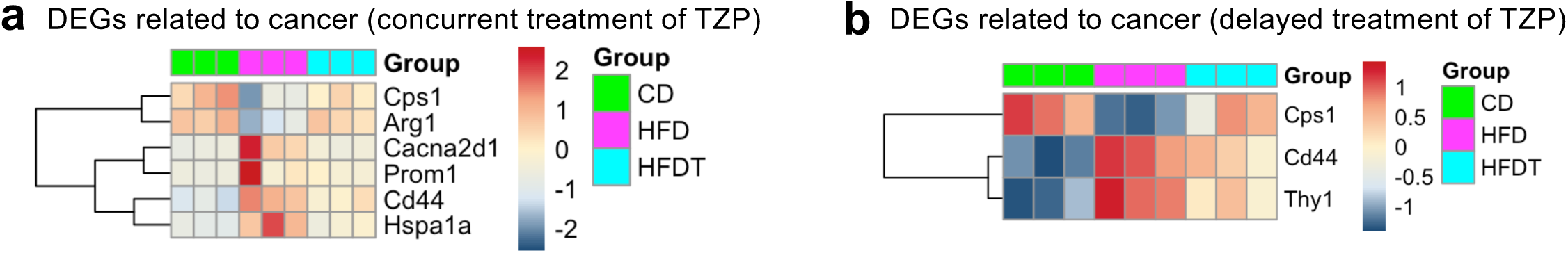
Both concurrent and delayed treatments of TZP reverse transcriptomic signatures related to cancer. **a,** Male C57BL/6J mice were fed CD or HFD with concurrent treatment of TZP or vehicle for 10 weeks (*n* = 3 per group). **b,** Male C57BL/6J mice were fed CD or HFD for 7 weeks, followed by treatment with TZP or vehicle for the subsequent 10 weeks along with the respective diets (*n* = 3 per group). **a,b,** Heatmap of cancer-related DEGs in the liver of mice.

**Extended Data Fig. 8:**
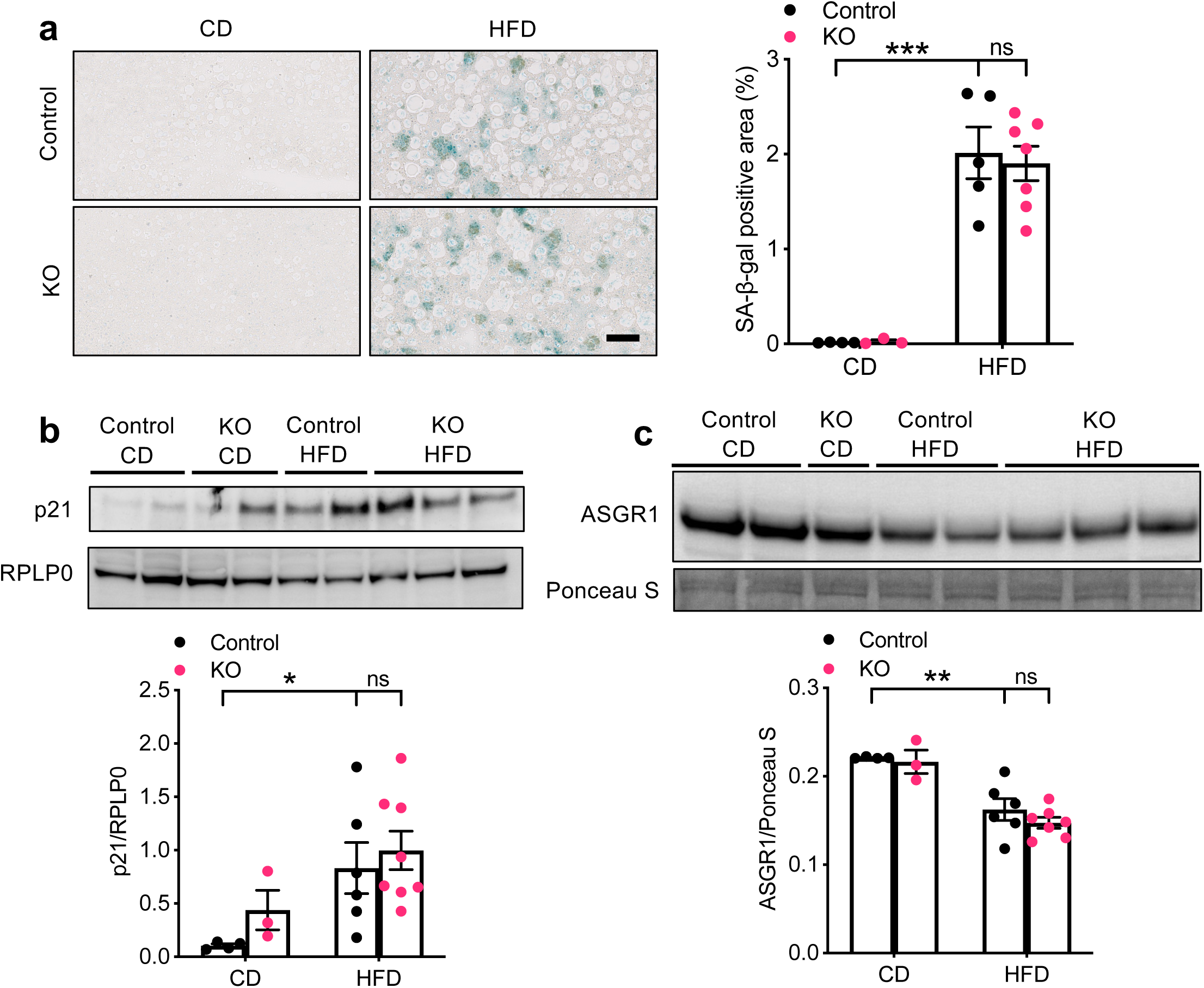
HFD-induced cellular senescence is not attenuated despite deficiency of NETosis. **a-c,** Male *Padi4^fl/fl^* (Control) and *Vav1-Cre Padi4^fl/fl^* (KO) were fed CD or HFD for 10 weeks. **a,** Representative images of SA-β-gal staining (left) and quantification of SA-β-gal positive area (%) (right) on the liver sections. Left to right on the bar chart, *n* = 4, 3, 5, 7. Scale bar, 100 µm. **b,** Representative Western blots and quantification of the levels of p21 (**b**) and ASGR1 (**c**). RPLP0 and Ponceau S-stained protein served as the loading controls. **b**, Left to right on the bar chart, *n* = 4, 3, 6, 8. **c,** Left to right on the bar chart, *n* = 4, 3, 6, 7. Statistical comparisons were made using two-tailed unpaired t-test. \*\*\**P* < 0.001, ns, not significant. Data are mean ± s.e.m.

**Extended Data Fig. 9:**
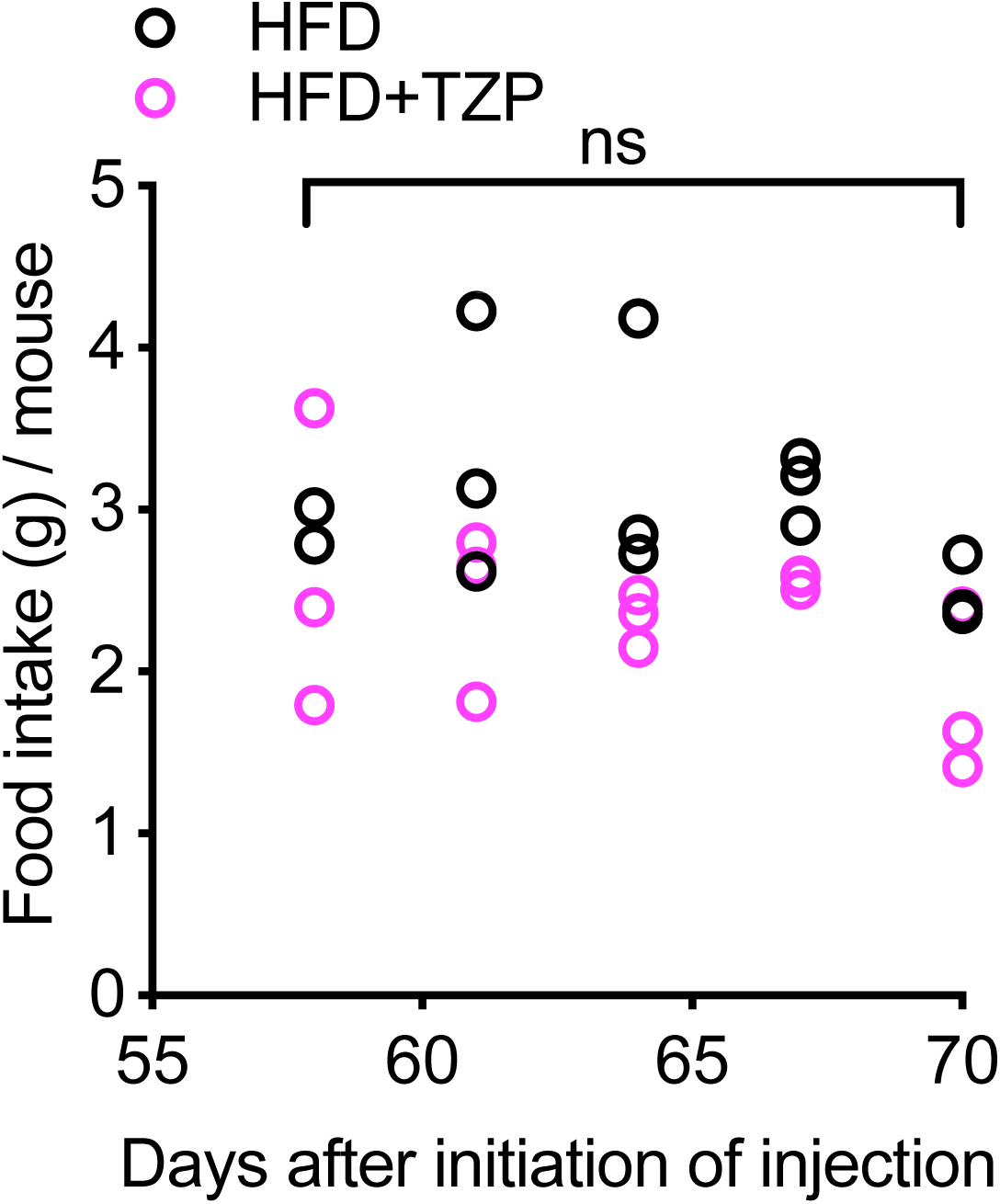
The TZP regimens do not affect food intake. Food intake (g) per mouse of HFD and HFD+TZP groups. Data was collected from 3 different batches of TZP treatments, with each batch containing 1 cage (3-5 mice per cage) per group. Each point was computed by dividing total food consumption of one cage of mice by the number of mice housed in that cage over 24 hours. Statistical comparisons were made using Mann-Whitney test. ns, not significant.

## References

1. Younossi, Z. M., Wong, G., Anstee, Q. M. & Henry, L. The Global Burden of Liver Disease. Clin. Gastroenterol. Hepatol. 21, 1978–1991 (2023).

2. Younossi, Z. M., Kalligeros, M. & Henry, L. Epidemiology of metabolic dysfunction-associated steatotic liver disease. Clin. Mol. Hepatol. 31, S32–S50 (2025).

3. Teng, M. L. et al. Global incidence and prevalence of nonalcoholic fatty liver disease. Clin. Mol. Hepatol. 29, S32–S42 (2023).

4. Stefan, N., Yki-Järvinen, H. & Neuschwander-Tetri, B. A. Metabolic dysfunction-associated steatotic liver disease: heterogeneous pathomechanisms and effectiveness of metabolism-based treatment. Lancet Diabetes Endocrinol. 13, 134–148 (2025).

5. Brenner, D. A. Reversibility of Liver Fibrosis. Gastroenterol. Hepatol. (NY) 9, 737–739 (2013).

6. Devarbhavi, H. et al. Global burden of liver disease: 2023 update. J. Hepatol. 79, 516–537 (2023).

7. Zheng, J. et al. Hepatocellular carcinoma: signaling pathways and therapeutic advances. Sig. Transduct. Target. Ther. 10, 1–43 (2025).

8. Saif, M. et al. Non-invasive monitoring of chronic liver disease via near-infrared and shortwave-infrared imaging of endogenous lipofuscin. Nat. Biomed. Eng. 4, 801–813 (2020).

9. Jia, H. et al. The role of altered lipid composition and distribution in liver fibrosis revealed by multimodal nonlinear optical microscopy. Sci. Adv. 9, eabq2937 (2023).

10. Luukkonen, P. K. et al. Saturated Fat Is More Metabolically Harmful for the Human Liver Than Unsaturated Fat or Simple Sugars. Diabetes Care 41, 1732–1739 (2018).

11. Chalasani, N. et al. Features and Outcomes of 899 Patients with Drug-induced Liver Injury: The DILIN Prospective Study. Gastroenterology 148, 1340–1352.e7 (2015).

12. Gray, D. A. & Woulfe, J. Lipofuscin and Aging: A Matter of Toxic Waste. Sci. Aging Knowl. Environ. 2005, re1 (2005).

13. American Association for the Study of Liver Diseases. Pathology Pearls: Evaluation of Donor Liver Biopsies. https://www.aasld.org/liver-fellow-network/core-series/pathology-pearls/pathology-pearls-evaluation-donor-liver-biopsies. Date of access: January 15, 2025.

14. Schmucker, D. L. & Sachs, H. Quantifying dense bodies and lipofuscin during aging: a morphologist’s perspective. Arch. Gerontol. Geriatr. 34, 249–261 (2002).

15. Pan, C. et al. Lipofuscin causes atypical necroptosis through lysosomal membrane permeabilization. Proc. Natl. Acad. Sci. U.S.A. 118, e2100122118 (2021).

16. Baldensperger, T. et al. The age pigment lipofuscin causes oxidative stress, lysosomal dysfunction, and pyroptotic cell death. Free Radic. Biol. Med. 225, 871–880 (2024).

17. Cho, Y. & Szabo, G. Two Faces of Neutrophils in Liver Disease Development and Progression. Hepatology 74, 503–512 (2021).

18. Liu, K., Wang, F.-S. & Xu, R. Neutrophils in liver diseases: pathogenesis and therapeutic targets. Cell. Mol. Immunol. 18, 38–44 (2021).

19. Brinkmann, V. et al. Neutrophil Extracellular Traps Kill Bacteria. Science 303, 1532–1535 (2004).

20. Papayannopoulos, V. Neutrophil extracellular traps in immunity and disease. Nat. Rev. Immunol. 18, 134–147 (2018).

21. Wong, S. L. et al. Diabetes primes neutrophils to undergo NETosis, which impairs wound healing. Nat. Med. 21, 815–819 (2015).

22. Fadini, G. P. et al. NETosis Delays Diabetic Wound Healing in Mice and Humans. Diabetes 65, 1061–1071 (2016).

23. Wang, L. et al. Hyperglycemia Induces Neutrophil Extracellular Traps Formation Through an NADPH Oxidase-Dependent Pathway in Diabetic Retinopathy. Front. Immunol. 9, 3076 (2019).

24. Li, T. et al. Cellular communication network factor 1 promotes retinal leakage in diabetic retinopathy via inducing neutrophil stasis and neutrophil extracellular traps extrusion. Cell Commun. Signal. 22, 275 (2024).

25. Zheng, F. et al. Neutrophil Extracellular Traps Induce Glomerular Endothelial Cell Dysfunction and Pyroptosis in Diabetic Kidney Disease. Diabetes 71, 2739–2750 (2022).

26. Van Bruggen, S. et al. Neutrophil peptidylarginine deiminase 4 is essential for detrimental age-related cardiac remodelling and dysfunction in mice. Phil. Trans. R. Soc. B 378, 20220475 (2023).

27. Suzuki, M. et al. PAD4 Deficiency Improves Bleomycin-induced Neutrophil Extracellular Traps and Fibrosis in Mouse Lung. Am. J. Respir. Cell Mol. Biol. 63, 806–818 (2020).

28. Ludvik, B. et al. Once-weekly tirzepatide versus once-daily insulin degludec as add-on to metformin with or without SGLT2 inhibitors in patients with type 2 diabetes (SURPASS-3): a randomised, open-label, parallel-group, phase 3 trial. The Lancet 398, 583–598 (2021).

29. Loomba, R. et al. Tirzepatide for Metabolic Dysfunction–Associated Steatohepatitis with Liver Fibrosis. NEJM 391, 299–310 (2024).

30. Matsumoto, M. et al. An improved mouse model that rapidly develops fibrosis in non-alcoholic steatohepatitis. Int. J. Exp. Path. 94, 93–103 (2013).

31. Puri, P. et al. A lipidomic analysis of nonalcoholic fatty liver disease. Hepatology 46, 1081–1090 (2007).

32. Terman, A. & Brunk, U. T. Lipofuscin. Int. J. Biochem. Cell Biol. 36, 1400–1404 (2004).

33. Sugasawa, T. et al. One Week of CDAHFD Induces Steatohepatitis and Mitochondrial Dysfunction with Oxidative Stress in Liver. Int. J. Mol. Sci. 22, 5851 (2021).

34. Evangelou, K. & Gorgoulis, V. G. Sudan Black B, The Specific Histochemical Stain for Lipofuscin: A Novel Method to Detect Senescent Cells. in Oncogene-Induced Senescence: Methods and Protocols 111–119 (Springer, New York, NY, 2017).

35. Escrevente, C. et al. Formation of Lipofuscin-Like Autofluorescent Granules in the Retinal Pigment Epithelium Requires Lysosome Dysfunction. Invest. Ophthalmol. Vis. Sci. 62, 39 (2021).

36. Ablonczy, Z. et al. The utilization of fluorescence to identify the components of lipofuscin by imaging mass spectrometry. Proteomics 14, 936–944 (2014).

37. Burns, J. C. et al. Differential accumulation of storage bodies with aging defines discrete subsets of microglia in the healthy brain. eLife 9, e57495 (2020).

38. Zhang, J. et al. Protective effect of autophagy on human retinal pigment epithelial cells against lipofuscin fluorophore A2E: implications for age-related macular degeneration. Cell Death Dis. 6, e1972–e1972 (2015).

39. Van Der Poorten, D. et al. Hepatic fat loss in advanced nonalcoholic steatohepatitis: Are alterations in serum adiponectin the cause? Hepatology 57, 2180–2188 (2013).

40. Bedossa, P. et al. Histopathological algorithm and scoring system for evaluation of liver lesions in morbidly obese patients. Hepatology 56, 1751–1759 (2012).

41. Wu, Y. et al. A Chemokine Receptor CXCR2 Macromolecular Complex Regulates Neutrophil Functions in Inflammatory Diseases. J. Biol. Chem. 287, 5744–5755 (2012).

42. Forsthuber, A. et al. CXCL5 as Regulator of Neutrophil Function in Cutaneous Melanoma. J. Invest. Dermatol. 139, 186–194 (2019).

43. Birnie, G. D. The HL60 cell line: a model system for studying human myeloid cell differentiation. Br. J. Cancer, Suppl. 9, 41–45 (1988).

44. Millius, A. & Weiner, O. D. Manipulation of Neutrophil-Like HL-60 Cells for the Study of Directed Cell Migration. in Live Cell Imaging 147–158 (Humana Press, 2010).

45. Thiam, H. R. et al. NETosis proceeds by cytoskeleton and endomembrane disassembly and PAD4-mediated chromatin decondensation and nuclear envelope rupture. Proc. Natl. Acad. Sci. U.S.A. 117, 7326–7337 (2020).

46. Wang, Y. et al. Histone hypercitrullination mediates chromatin decondensation and neutrophil extracellular trap formation. J. Cell Biol. 184, 205–213 (2009).

47. Yamada, K. et al. Characteristics of hepatic fatty acid compositions in patients with nonalcoholic steatohepatitis. Liver Int. 35, 582–590 (2015).

48. Münzer, P. et al. NLRP3 Inflammasome Assembly in Neutrophils Is Supported by PAD4 and Promotes NETosis Under Sterile Conditions. Front. Immunol. 12, 683803 (2021).

49. Li, W. et al. Reducing lipofuscin accumulation and cardiomyocytic senescence of aging heart by enhancing autophagy. Exp. Cell Res. 403, 112585 (2021).

50. Aravinthan, A. D. & Alexander, G. J. M. Senescence in chronic liver disease: Is the future in aging? J. Hepatol. 65, 825–834 (2016).

51. Kim, H., Kisseleva, T. & Brenner, D. A. Aging and liver disease. Curr. Opin. Gastroenterol. 31, 184–191 (2015).

52. Ogrodnik, M. et al. Cellular senescence drives age-dependent hepatic steatosis. Nat. Commun. 8, 15691 (2017).

53. Schaum, N. et al. Ageing hallmarks exhibit organ-specific temporal signatures. Nature 583, 596–602 (2020).

54. Saul, D. et al. A new gene set identifies senescent cells and predicts senescence-associated pathways across tissues. Nat. Commun. 13, 4827 (2022).

55. Ogrodnik, M. et al. Guidelines for minimal information on cellular senescence experimentation in vivo. Cell 187, 4150–4175 (2024).

56. Yang, J.-H. et al. Loss of epigenetic information as a cause of mammalian aging. Cell 186, 305–326.e27 (2023).

57. Wang, M.-J., Chen, F., Lau, J. T. Y. & Hu, Y.-P. Hepatocyte polyploidization and its association with pathophysiological processes. Cell Death Dis. 8, e2805–e2805 (2017).

58. Wuestefeld, A. et al. A Pro-Regenerative Environment Triggers Premalignant to Malignant Transformation of Senescent Hepatocytes. Cancer Research 83, 428–440 (2023).

59. Mansouri, V. et al. Assessment of liver cancer biomarkers. Gastroenterol. Hepatol. Bed Bench 13, S29–S39 (2020).

60. Nio, K., Yamashita, T. & Kaneko, S. The evolving concept of liver cancer stem cells. Mol. Cancer 16, 4 (2017).

61. Brunk, U. T. & Terman, A. Lipofuscin: mechanisms of age-related accumulation and influence on cell function. Free Radic. Biol. Med. 33, 611–619 (2002).

62. Brandstetter, C., Mohr, L. K. M., Latz, E., Holz, F. G. & Krohne, T. U. Light induces NLRP3 inflammasome activation in retinal pigment epithelial cells via lipofuscin-mediated photooxidative damage. J. Mol. Med. (Berl) 93, 905–916 (2015).

63. Anderson, O. A., Finkelstein, A. & Shima, D. T. A2E Induces IL-1ß Production in Retinal Pigment Epithelial Cells via the NLRP3 Inflammasome. PLoS One 8, e67263 (2013).

64. Walter, S. et al. Oxidized protein aggregate lipofuscin impairs cardiomyocyte contractility via late-stage autophagy inhibition. Redox Biol. 81, 103559 (2025).

65. Xia, Y. et al. Neutrophil extracellular traps promote MASH fibrosis by metabolic reprogramming of HSC. Hepatology 81, 947–961 (2024).

66. Kolaczkowska, E. et al. Molecular mechanisms of NET formation and degradation revealed by intravital imaging in the liver vasculature. Nat. Commun. 6, 6673 (2015).

67. Ying, Z. et al. Combined GIP receptor and GLP1 receptor agonism attenuates NAFLD in male APOE∗3-Leiden.CETP mice. eBioMedicine 93, 104684 (2023).

68. Regmi, A. et al. Tirzepatide modulates the regulation of adipocyte nutrient metabolism through long-acting activation of the GIP receptor. Cell Metab. 36, 1534–1549.e7 (2024).

69. Ravussin, E. et al. Tirzepatide did not impact metabolic adaptation in people with obesity, but increased fat oxidation. Cell Metab. 37, 1060–1074.e4 (2025).

70. Höhn, A. & Grune, T. Lipofuscin: formation, effects and role of macroautophagy. Redox Biol. 1, 140–144 (2013).

71. Aravinthan, A. et al. Hepatocyte senescence predicts progression in non-alcohol-related fatty liver disease. J. Hepatol. 58, 549–556 (2013).

72. Krizhanovsky, V. et al. Senescence of Activated Stellate Cells Limits Liver Fibrosis. Cell 134, 657–667 (2008).

73. Grosse, L. et al. Defined p16High Senescent Cell Types Are Indispensable for Mouse Healthspan. Cell Metab. 32, 87–99.e6 (2020).

74. Umbaugh, D. S., Nguyen, N. T., Smith, S. H., Ramachandran, A. & Jaeschke, H. The p21+ perinecrotic hepatocytes produce the chemokine CXCL14 after a severe acetaminophen overdose promoting hepatocyte injury and delaying regeneration. Toxicology 504, 153804 (2024).

75. Raffaele, M. et al. Mild exacerbation of obesity- and age-dependent liver disease progression by senolytic cocktail dasatinib + quercetin. Cell Commun. Signal. 19, 44 (2021).

76. Sagy, Y. W. et al. Glucagon-like peptide-1 receptor agonists compared with bariatric metabolic surgery and the risk of obesity-related cancer: an observational, retrospective cohort study. eClinicalMedicine 83, 103213 (2025).

77. Coskun, T. et al. LY3298176, a novel dual GIP and GLP-1 receptor agonist for the treatment of type 2 diabetes mellitus: From discovery to clinical proof of concept. Mol. Metab. 18, 3–14 (2018).

78. Samms, R. J. et al. Tirzepatide induces a thermogenic-like amino acid signature in brown adipose tissue. Mol. Metab. 64, 101550 (2022).

79. Urva, S. et al. Effects of Hepatic Impairment on the Pharmacokinetics of the Dual GIP and GLP-1 Receptor Agonist Tirzepatide. Clin. Pharmacokinet. 61, 1057–1067 (2022).

80. Mehlem, A., Hagberg, C. E., Muhl, L., Eriksson, U. & Falkevall, A. Imaging of neutral lipids by oil red O for analyzing the metabolic status in health and disease. Nat. Protoc. 8, 1149–1154 (2013).

81. Kim, D., Paggi, J. M., Park, C., Bennett, C. & Salzberg, S. L. Graph-Based Genome Alignment and Genotyping with HISAT2 and HISAT-genotype. Nat. Biotechnol. 37, 907–915 (2019).

82. Liao, Y., Smyth, G. K. & Shi, W. featureCounts: an efficient general purpose program for assigning sequence reads to genomic features. Bioinformatics 30, 923–930 (2014).

83. Love, M. I., Huber, W. & Anders, S. Moderated estimation of fold change and dispersion for RNA-seq data with DESeq2. Genome Biol. 15, 550 (2014).

84. Yu, G., Wang, L.-G., Han, Y. & He, Q.-Y. clusterProfiler: an R Package for Comparing Biological Themes Among Gene Clusters. OMICS 16, 284–287 (2012).

85. Gustavsson, E. K., Zhang, D., Reynolds, R. H., Garcia-Ruiz, S. & Ryten, M. ggtranscript: an R package for the visualization and interpretation of transcript isoforms using ggplot2. Bioinformatics 38, 3844–3846 (2022).

