## Supplementary Tables for "Tirzepatide reduces diet-induced senescence and NETosis-mediated liver fibrosis in mice"

**Supplementary Table 1: DEGs in the fibrosis-related pathways in the liver of mice with concurrent treatment of TZP along with the commencement of HFD (HFD+TZP versus HFD)**

| gene_id | external_gene_name | gene_biotype | log2FoldChange | pvalue | padj |
|---|---|---|---|---|---|
| ENSMUSG00000000049 | Apoh | protein_coding | 0.75237549 | 0.005528756 | 0.043917889 |
| ENSMUSG00000000058 | Cav2 | protein_coding | -0.649339183 | 0.003497412 | 0.03191097 |
| ENSMUSG00000000244 | Tspan32 | protein_coding | -0.990385933 | 0.002252039 | 0.023509881 |
| ENSMUSG00000000489 | Pdgfb | protein_coding | -1.113454629 | 0.000190482 | 0.003666029 |
| ENSMUSG00000001119 | Col6a1 | protein_coding | -1.164276809 | 0.000758902 | 0.010653028 |
| ENSMUSG00000001131 | Timp1 | protein_coding | -2.077632697 | 4.57555E-10 | 6.93505E-08 |
| ENSMUSG00000001506 | Col1a1 | protein_coding | -2.212354291 | 1.49461E-09 | 1.96951E-07 |
| ENSMUSG00000001555 | Fkbp10 | protein_coding | -1.198176759 | 0.000113747 | 0.002532387 |
| ENSMUSG00000001870 | Ltbp1 | protein_coding | -1.117459702 | 4.15152E-08 | 3.68026E-06 |
| ENSMUSG00000002111 | Spi1 | protein_coding | -0.722506383 | 5.67756E-05 | 0.001496487 |
| ENSMUSG00000002602 | Axl | protein_coding | -0.671745606 | 0.00502282 | 0.041344285 |
| ENSMUSG00000002900 | Lamb1 | protein_coding | -0.625581975 | 0.005188692 | 0.042114169 |
| ENSMUSG00000003534 | Ddr1 | protein_coding | -1.917593196 | 5.07474E-05 | 0.001376839 |
| ENSMUSG00000004266 | Ptpn6 | protein_coding | -0.599475983 | 0.001006912 | 0.012975969 |
| ENSMUSG00000005087 | Cd44 | protein_coding | -0.962391743 | 3.72252E-05 | 0.00108249 |
| ENSMUSG00000005124 | Ccn4 | protein_coding | -1.658038022 | 1.59048E-05 | 0.000536037 |
| ENSMUSG00000006205 | Htra1 | protein_coding | -2.357393317 | 1.30105E-05 | 0.000456521 |
| ENSMUSG00000006369 | Fbln1 | protein_coding | -1.799153007 | 0.003374639 | 0.031211292 |
| ENSMUSG00000006403 | Adamts4 | protein_coding | -1.93928207 | 0.0001508 | 0.003124295 |
| ENSMUSG00000007613 | Tgfbr1 | protein_coding | -0.608097892 | 0.003801824 | 0.033866032 |
| ENSMUSG00000015085 | Entpd2 | protein_coding | -0.731014695 | 0.002110643 | 0.022518149 |
| ENSMUSG00000015850 | Adamtsl4 | protein_coding | 0.753921851 | 0.003756205 | 0.033610637 |
| ENSMUSG00000016028 | Celsr1 | protein_coding | 1.449093202 | 0.000187836 | 0.003624635 |
| ENSMUSG00000016458 | Wt1 | protein_coding | -1.649367703 | 0.000185224 | 0.003593118 |
| ENSMUSG00000016494 | Cd34 | protein_coding | -1.342576784 | 0.003335468 | 0.031025757 |
| ENSMUSG00000016918 | Sulf1 | protein_coding | -2.005218814 | 1.6928E-07 | 1.23803E-05 |
| ENSMUSG00000017950 | Hnf4a | protein_coding | 0.538909077 | 0.00296725 | 0.02846031 |
| ENSMUSG00000019539 | Rcn3 | protein_coding | -1.12021333 | 1.61316E-05 | 0.000542429 |
| ENSMUSG00000019726 | Lyst | protein_coding | 0.864069996 | 5.41457E-06 | 0.000228239 |
| ENSMUSG00000019899 | Lama2 | protein_coding | -0.961509656 | 0.002161794 | 0.022913454 |
| ENSMUSG00000019987 | Arg1 | protein_coding | 1.153509948 | 1.66857E-08 | 1.62708E-06 |
| ENSMUSG00000020120 | Plek | protein_coding | -1.055749136 | 0.000142577 | 0.003005004 |
| ENSMUSG00000020473 | Aebp1 | protein_coding | -1.328975834 | 0.001667415 | 0.018985864 |
| ENSMUSG00000020810 | Cygb | protein_coding | -1.215990346 | 4.46634E-05 | 0.001246739 |
| ENSMUSG00000020900 | Myh10 | protein_coding | -1.65281503 | 2.68384E-15 | 9.3468E-13 |
| ENSMUSG00000021136 | Smoc1 | protein_coding | 1.13999083 | 6.1506E-16 | 2.49902E-13 |
| ENSMUSG00000021253 | Tgfb3 | protein_coding | -1.488947807 | 0.000506464 | 0.007872525 |
| ENSMUSG00000021262 | Evl | protein_coding | -0.917393568 | 0.000515444 | 0.007992521 |
| ENSMUSG00000021457 | Syk | protein_coding | -0.835136176 | 0.000127213 | 0.002756315 |
| ENSMUSG00000021492 | F12 | protein_coding | 0.53665752 | 0.005325899 | 0.042795626 |
| ENSMUSG00000021822 | Plau | protein_coding | -0.745168049 | 0.001785224 | 0.019887644 |
| ENSMUSG00000021944 | Gata4 | protein_coding | 0.466564218 | 0.002991238 | 0.028671589 |
| ENSMUSG00000021994 | Wnt5a | protein_coding | -0.956559381 | 0.003855549 | 0.034116824 |
| ENSMUSG00000021998 | Lcp1 | protein_coding | -0.699208923 | 0.000125725 | 0.002732514 |
| ENSMUSG00000022010 | Tsc22d1 | protein_coding | -0.768803512 | 9.67088E-05 | 0.002238226 |
| ENSMUSG00000022371 | Col14a1 | protein_coding | -0.814279692 | 0.000135998 | 0.002904006 |
| ENSMUSG00000022512 | Cldn1 | protein_coding | 0.681142995 | 0.001475601 | 0.017280721 |
| ENSMUSG00000022665 | Ccdc80 | protein_coding | -1.538753531 | 9.43987E-06 | 0.000355869 |
| ENSMUSG00000022780 | Meltf | protein_coding | 2.232401756 | 0.000238212 | 0.004424625 |
| ENSMUSG00000022817 | Itgb5 | protein_coding | -0.743608789 | 5.2289E-05 | 0.001408521 |
| ENSMUSG00000022875 | Kng1 | protein_coding | 0.678797396 | 0.000111664 | 0.002497419 |
| ENSMUSG00000022894 | Adamts5 | protein_coding | -1.549648011 | 6.81007E-05 | 0.001716771 |
| ENSMUSG00000023067 | Cdkn1a | protein_coding | -1.246863973 | 0.000336958 | 0.005801602 |
| ENSMUSG00000023191 | P3h3 | protein_coding | -1.100007604 | 0.001171786 | 0.014675556 |
| ENSMUSG00000023224 | Serping1 | protein_coding | -0.584185078 | 0.000659598 | 0.009614739 |
| ENSMUSG00000023886 | Smoc2 | protein_coding | -0.921781385 | 0.001179489 | 0.014707912 |
| ENSMUSG00000023972 | Ptk7 | protein_coding | -1.300718992 | 0.001317274 | 0.015897491 |
| ENSMUSG00000024053 | Emilin2 | protein_coding | -0.735427369 | 0.000922225 | 0.0121526 |
| ENSMUSG00000024087 | Cyp1b1 | protein_coding | -1.801590023 | 1.48338E-05 | 0.000506695 |

| gene_id | external_gene_name | gene_biotype | log2FoldChange | pvalue | padj |
|---|---|---|---|---|---|
| ENSMUSG00000024299 | Adamts10 | protein_coding | -1.104207899 | 0.000469819 | 0.007461975 |
| ENSMUSG00000024529 | Lox | protein_coding | -2.434477177 | 3.23866E-12 | 7.8953E-10 |
| ENSMUSG00000024598 | Fbn2 | protein_coding | -4.137890403 | 6.74464E-05 | 0.001704509 |
| ENSMUSG00000024610 | Cd74 | protein_coding | -0.936314374 | 0.001194643 | 0.014858877 |
| ENSMUSG00000024620 | Pdgfrb | protein_coding | -1.225770534 | 3.64254E-05 | 0.001065587 |
| ENSMUSG00000024659 | Anxa1 | protein_coding | -1.819659609 | 6.50892E-11 | 1.19007E-08 |
| ENSMUSG00000024696 | Lpxn | protein_coding | -1.227913062 | 1.82089E-05 | 0.000599869 |
| ENSMUSG00000024909 | Efemp2 | protein_coding | -1.147924723 | 3.86556E-05 | 0.001110836 |
| ENSMUSG00000024940 | Ltbp3 | protein_coding | -1.817063537 | 5.58289E-08 | 4.86077E-06 |
| ENSMUSG00000024965 | Fermt3 | protein_coding | -0.780683161 | 0.000611455 | 0.009089174 |
| ENSMUSG00000025225 | Nfkb2 | protein_coding | -0.903385684 | 0.000288028 | 0.005102948 |
| ENSMUSG00000025321 | Itgb8 | protein_coding | -2.045327326 | 0.000164084 | 0.003324173 |
| ENSMUSG00000025856 | Pdgfa | protein_coding | -0.89328765 | 0.000242847 | 0.004490674 |
| ENSMUSG00000025880 | Smad7 | protein_coding | 0.909195117 | 0.000676003 | 0.009789998 |
| ENSMUSG00000026042 | Col5a2 | protein_coding | -2.01753336 | 1.80671E-07 | 1.31476E-05 |
| ENSMUSG00000026043 | Col3a1 | protein_coding | -1.724410946 | 1.32934E-07 | 1.01109E-05 |
| ENSMUSG00000026177 | Slc11a1 | protein_coding | -0.992657611 | 2.07404E-05 | 0.000671173 |
| ENSMUSG00000026574 | Dpt | protein_coding | -2.454880488 | 1.82746E-08 | 1.77022E-06 |
| ENSMUSG00000026580 | Selp | protein_coding | 0.916972857 | 0.005063433 | 0.041538327 |
| ENSMUSG00000026715 | Serpinc1 | protein_coding | 0.71100032 | 0.001718217 | 0.019302892 |
| ENSMUSG00000026728 | Vim | protein_coding | -1.143970369 | 6.47227E-05 | 0.001657967 |
| ENSMUSG00000026768 | Itga8 | protein_coding | -2.519310983 | 2.26139E-09 | 2.82264E-07 |
| ENSMUSG00000026837 | Col5a1 | protein_coding | -1.068806459 | 0.000134748 | 0.002885895 |
| ENSMUSG00000026971 | Itgb6 | protein_coding | -3.115583526 | 0.001866049 | 0.020568723 |
| ENSMUSG00000027087 | Itgav | protein_coding | -0.63786422 | 0.005261677 | 0.042403605 |
| ENSMUSG00000027204 | Fbn1 | protein_coding | -1.577680355 | 1.90291E-05 | 0.000624079 |
| ENSMUSG00000027230 | Creb3l1 | protein_coding | -0.991776921 | 0.00015692 | 0.003210167 |
| ENSMUSG00000027315 | Spint1 | protein_coding | -2.081276953 | 0.000503147 | 0.007846423 |
| ENSMUSG00000027611 | Procr | protein_coding | -0.898482472 | 0.000546261 | 0.00834917 |
| ENSMUSG00000027646 | Src | protein_coding | -1.007148866 | 0.001405063 | 0.016681701 |
| ENSMUSG00000027712 | Anxa5 | protein_coding | -1.068742462 | 6.86305E-10 | 9.7462E-08 |
| ENSMUSG00000027831 | Veph1 | protein_coding | -2.338343761 | 1.1064E-07 | 8.65415E-06 |
| ENSMUSG00000028249 | Sdcbp | protein_coding | -0.699877052 | 0.000124044 | 0.002704014 |
| ENSMUSG00000028339 | Col15a1 | protein_coding | -2.953131769 | 2.2771E-09 | 2.82264E-07 |
| ENSMUSG00000028369 | Svep1 | protein_coding | -2.977473343 | 3.63273E-11 | 7.18053E-09 |
| ENSMUSG00000028420 | Tmem38b | protein_coding | 0.626276293 | 0.002738306 | 0.026941301 |
| ENSMUSG00000028599 | Tnfrsf1b | protein_coding | -0.46629188 | 0.006516057 | 0.04933249 |
| ENSMUSG00000028664 | Ephb2 | protein_coding | -3.175520133 | 9.72116E-16 | 3.74188E-13 |
| ENSMUSG00000029061 | Mmp23 | protein_coding | -1.267402687 | 0.001023707 | 0.01314864 |
| ENSMUSG00000029163 | Emilin1 | protein_coding | -0.864280945 | 0.000406904 | 0.006718022 |
| ENSMUSG00000029185 | Fam114a1 | protein_coding | -0.706574293 | 1.08846E-05 | 0.000396044 |
| ENSMUSG00000029377 | Ereg | protein_coding | -1.314223196 | 3.0564E-05 | 0.000927325 |
| ENSMUSG00000029581 | Fscn1 | protein_coding | -0.646859702 | 0.004672924 | 0.039463548 |
| ENSMUSG00000029661 | Col1a2 | protein_coding | -1.78764272 | 9.43696E-09 | 9.78967E-07 |
| ENSMUSG00000029675 | Eln | protein_coding | -1.564547529 | 2.39013E-05 | 0.00076001 |
| ENSMUSG00000029913 | Prdm5 | protein_coding | -2.600411625 | 0.00229255 | 0.023698321 |
| ENSMUSG00000030022 | Adamts9 | protein_coding | -1.170810317 | 0.000950687 | 0.012404733 |
| ENSMUSG00000030342 | Cd9 | protein_coding | -1.272995446 | 2.31054E-05 | 0.000739525 |
| ENSMUSG00000030530 | Furin | protein_coding | 0.712413655 | 2.43087E-05 | 0.000763012 |
| ENSMUSG00000030774 | Pak1 | protein_coding | -0.814595848 | 0.002579554 | 0.025772637 |
| ENSMUSG00000031328 | Flna | protein_coding | -0.807669742 | 0.00136835 | 0.01636538 |
| ENSMUSG00000031342 | Gpm6b | protein_coding | -1.457561089 | 0.001256037 | 0.015408833 |
| ENSMUSG00000031380 | Vegfd | protein_coding | -1.135195204 | 0.005671343 | 0.044599319 |
| ENSMUSG00000031453 | Rasa3 | protein_coding | -0.622237589 | 0.003804053 | 0.033866032 |
| ENSMUSG00000031502 | Col4a1 | protein_coding | -0.832274966 | 0.001092876 | 0.013808682 |
| ENSMUSG00000031538 | Plat | protein_coding | -2.231223877 | 3.62158E-08 | 3.33162E-06 |
| ENSMUSG00000031740 | Mmp2 | protein_coding | -1.93651247 | 2.31905E-07 | 1.65467E-05 |
| ENSMUSG00000031778 | Cx3cl1 | protein_coding | -2.783479628 | 3.75672E-08 | 3.41301E-06 |
| ENSMUSG00000031790 | Mmp15 | protein_coding | -0.832513131 | 0.002524879 | 0.025417344 |
| ENSMUSG00000031825 | Crispld2 | protein_coding | -1.974324946 | 2.19311E-05 | 0.000705026 |
| ENSMUSG00000031980 | Agt | protein_coding | 1.323350059 | 3.11871E-18 | 1.75451E-15 |
| ENSMUSG00000032068 | Plet1 | protein_coding | -1.408759155 | 0.000841441 | 0.011417218 |

| gene_id | external_gene_name | gene_biotype | log2FoldChange | pvalue | padj |
|---|---|---|---|---|---|
| ENSMUSG00000032231 | Anxa2 | protein_coding | -1.22194755 | 3.96088E-06 | 0.000174506 |
| ENSMUSG00000032334 | Loxl1 | protein_coding | -2.217016228 | 2.9692E-07 | 2.05832E-05 |
| ENSMUSG00000032374 | Plod2 | protein_coding | -1.549532204 | 5.32261E-05 | 0.001419896 |
| ENSMUSG00000032382 | Snx1 | protein_coding | -0.483619095 | 0.0037496 | 0.033606248 |
| ENSMUSG00000033420 | Antxr1 | protein_coding | -1.6688623 | 3.97321E-05 | 0.001132869 |
| ENSMUSG00000034205 | Loxl2 | protein_coding | -1.33485325 | 1.66141E-07 | 1.22118E-05 |
| ENSMUSG00000034652 | Cd300a | protein_coding | -0.86480029 | 0.000544072 | 0.008324421 |
| ENSMUSG00000035270 | Impg2 | protein_coding | 1.423347273 | 0.000228904 | 0.004281552 |
| ENSMUSG00000035273 | Hpse | protein_coding | -0.688247082 | 0.003503727 | 0.03191097 |
| ENSMUSG00000035493 | Tgfbi | protein_coding | -0.514363287 | 1.12502E-05 | 0.000406312 |
| ENSMUSG00000036040 | Adamtsl2 | protein_coding | -1.903473612 | 2.22392E-09 | 2.80425E-07 |
| ENSMUSG00000036545 | Adamts2 | protein_coding | -1.38379537 | 6.79528E-07 | 4.1065E-05 |
| ENSMUSG00000036894 | Rap2b | protein_coding | -0.941029859 | 0.000238973 | 0.004424625 |
| ENSMUSG00000038400 | Pmepa1 | protein_coding | -1.511138655 | 0.000178536 | 0.003517339 |
| ENSMUSG00000038642 | Ctss | protein_coding | -1.328955945 | 2.56851E-07 | 1.80623E-05 |
| ENSMUSG00000039087 | Rreb1 | protein_coding | 0.402676559 | 0.0062227 | 0.047822086 |
| ENSMUSG00000039239 | Tgfb2 | protein_coding | -1.598063103 | 0.00070217 | 0.010069253 |
| ENSMUSG00000039546 | Ajap1 | protein_coding | -3.729658248 | 7.4364E-09 | 7.93957E-07 |
| ENSMUSG00000039942 | Ptger4 | protein_coding | -1.46024729 | 3.31663E-05 | 0.000990047 |
| ENSMUSG00000040152 | Thbs1 | protein_coding | -1.441303652 | 3.39046E-05 | 0.001005927 |
| ENSMUSG00000040690 | Col16a1 | protein_coding | -1.145401544 | 0.000237777 | 0.004424625 |
| ENSMUSG00000040711 | Sh3pxd2b | protein_coding | -1.315868956 | 0.000464896 | 0.007415524 |
| ENSMUSG00000041959 | S100a10 | protein_coding | -0.653524037 | 2.78222E-05 | 0.000858556 |
| ENSMUSG00000042613 | Pbxip1 | protein_coding | -0.586743432 | 0.006468109 | 0.049135271 |
| ENSMUSG00000043903 | Zfp469 | protein_coding | -2.930397446 | 0.000628441 | 0.009322721 |
| ENSMUSG00000045382 | Cxcr4 | protein_coding | -1.742667139 | 3.72418E-09 | 4.28925E-07 |
| ENSMUSG00000045672 | Col27a1 | protein_coding | -1.468747434 | 6.16658E-05 | 0.001588004 |
| ENSMUSG00000045991 | Onecut2 | protein_coding | 0.476675085 | 0.006282917 | 0.048095914 |
| ENSMUSG00000047407 | Tgif1 | protein_coding | -0.934994386 | 0.000104663 | 0.002366166 |
| ENSMUSG00000047414 | Flrt2 | protein_coding | -1.463603945 | 0.000417038 | 0.006846264 |
| ENSMUSG00000047497 | Adamts12 | protein_coding | -1.927261215 | 2.91084E-07 | 2.02747E-05 |
| ENSMUSG00000048612 | Myof | protein_coding | -1.494858538 | 8.26149E-08 | 6.67629E-06 |
| ENSMUSG00000049103 | Ccr2 | protein_coding | -0.859202799 | 0.003759736 | 0.033610637 |
| ENSMUSG00000049538 | Adamts16 | protein_coding | -4.068668753 | 0.000744731 | 0.010514646 |
| ENSMUSG00000049723 | Mmp12 | protein_coding | -2.064863543 | 2.21971E-09 | 2.80425E-07 |
| ENSMUSG00000050335 | Lgals3 | protein_coding | -1.335391538 | 1.50086E-07 | 1.1258E-05 |
| ENSMUSG00000050989 | Selenon | protein_coding | -1.123844536 | 0.000292388 | 0.005158936 |
| ENSMUSG00000052384 | Nrros | protein_coding | -0.747656147 | 5.20295E-06 | 0.000219952 |
| ENSMUSG00000054342 | Kcnn4 | protein_coding | -1.952277177 | 3.42941E-07 | 2.34402E-05 |
| ENSMUSG00000056220 | Pla2g4a | protein_coding | -0.754372584 | 0.003533427 | 0.032101513 |
| ENSMUSG00000058715 | Fcer1g | protein_coding | -0.953490669 | 0.000682474 | 0.009843691 |
| ENSMUSG00000060459 | Kng2 | protein_coding | 0.712752959 | 6.8192E-05 | 0.001716771 |
| ENSMUSG00000061048 | Cdh3 | protein_coding | -2.299854196 | 3.20563E-08 | 2.98655E-06 |
| ENSMUSG00000061878 | Sphk1 | protein_coding | -1.794501049 | 0.000274328 | 0.0049061 |
| ENSMUSG00000061947 | Serpina10 | protein_coding | -0.533203814 | 0.002996778 | 0.028687091 |
| ENSMUSG00000062960 | Kdr | protein_coding | 0.779128054 | 0.002011153 | 0.021623277 |
| ENSMUSG00000066113 | Adamtsl1 | protein_coding | -1.550829729 | 0.002010724 | 0.021623277 |
| ENSMUSG00000068196 | Col8a1 | protein_coding | -1.616185376 | 0.001500564 | 0.017502989 |
| ENSMUSG00000068923 | Syt11 | protein_coding | -1.252301959 | 0.00304107 | 0.029035069 |
| ENSMUSG00000070323 | Mmp27 | protein_coding | -1.366717778 | 0.001052693 | 0.013447809 |
| ENSMUSG00000070436 | Serpinh1 | protein_coding | -1.494093332 | 2.60755E-06 | 0.000124642 |
| ENSMUSG00000074272 | Ceacam1 | protein_coding | 0.604367664 | 0.005183389 | 0.042114169 |
| ENSMUSG00000079293 | Clec7a | protein_coding | -0.9486647 | 3.81682E-06 | 0.000169692 |
| ENSMUSG00000082361 | Btc | protein_coding | -1.634039675 | 6.09267E-08 | 5.15017E-06 |
| ENSMUSG00000112148 | Lilrb4a | protein_coding | -1.386049472 | 0.000148737 | 0.003099107 |

**Supplementary Table 2: DEGs in the fibrosis-related pathways in the liver of mice with delayed treatment of TZP starting 7 weeks post-HFD (HFD+TZP versus HFD)**

| gene_id | external_gene_name | gene_biotype | log2FoldChange | pvalue | padj |
|---|---|---|---|---|---|
| ENSMUSG00000000489 | Pdgfb | protein_coding | -0.895040654 | 0.000970402 | 0.018097205 |
| ENSMUSG00000000530 | Acvrl1 | protein_coding | -0.928574436 | 2.1694072759785e-05 | 0.001305324 |
| ENSMUSG00000001131 | Timp1 | protein_coding | -2.00425517 | 6.53050920364277e-08 | 1.6157833429395e-05 |
| ENSMUSG00000001506 | Col1a1 | protein_coding | -1.739881608 | 4.21603913982719e-07 | 6.57790642886532e-05 |
| ENSMUSG00000002111 | Spi1 | protein_coding | -0.874385987 | 3.49768455784642e-05 | 0.001873175 |
| ENSMUSG00000002233 | Rhoc | protein_coding | -0.863899618 | 0.001965869 | 0.030104906 |
| ENSMUSG00000002384 | Bmp8b | protein_coding | -2.884390272 | 0.000906162 | 0.017199057 |
| ENSMUSG00000002603 | Tgfb1 | protein_coding | -0.569169539 | 0.002852038 | 0.038548695 |
| ENSMUSG00000003051 | Elf3 | protein_coding | -1.14649724 | 0.002376349 | 0.033889697 |
| ENSMUSG00000003873 | Bax | protein_coding | -0.672952591 | 0.000265782 | 0.007651077 |
| ENSMUSG00000005087 | Cd44 | protein_coding | -0.694571184 | 0.002221897 | 0.032474934 |
| ENSMUSG00000006205 | Htra1 | protein_coding | -1.619718105 | 7.9697711250323e-07 | 0.000101898 |
| ENSMUSG00000006651 | Aplp1 | protein_coding | -1.125312004 | 0.002746635 | 0.037634462 |
| ENSMUSG00000015085 | Entpd2 | protein_coding | -1.403560471 | 6.86370798540654e-06 | 0.000541165 |
| ENSMUSG00000016028 | Celsr1 | protein_coding | 1.750683958 | 1.46041089379384e-06 | 0.000161923 |
| ENSMUSG00000017776 | Crk | protein_coding | 0.424367425 | 0.003774787 | 0.045669807 |
| ENSMUSG00000019539 | Rcn3 | protein_coding | -0.988052696 | 2.76065977400853e-06 | 0.000266526 |
| ENSMUSG00000019942 | Cdk1 | protein_coding | -1.050537388 | 0.000120533 | 0.004377186 |
| ENSMUSG00000020072 | Pbld2 | protein_coding | 0.836830326 | 0.003329574 | 0.042303774 |
| ENSMUSG00000020120 | Plek | protein_coding | -0.777301082 | 0.00158461 | 0.025840807 |
| ENSMUSG00000021223 | Papln | protein_coding | 1.307924102 | 0.000314674 | 0.008621332 |
| ENSMUSG00000021262 | Evl | protein_coding | -0.835106553 | 0.000730314 | 0.014934605 |
| ENSMUSG00000021822 | Plau | protein_coding | -0.867888911 | 0.001700028 | 0.027108763 |
| ENSMUSG00000021831 | Ero1a | protein_coding | 0.573781377 | 0.00321263 | 0.041298594 |
| ENSMUSG00000021948 | Prkcd | protein_coding | -0.68574173 | 0.002160414 | 0.031904889 |
| ENSMUSG00000022383 | Ppara | protein_coding | 1.099254378 | 1.55266407254882e-07 | 3.14791550251613e-05 |
| ENSMUSG00000022607 | Ptk2 | protein_coding | 0.496717372 | 0.003513502 | 0.043913761 |
| ENSMUSG00000022665 | Ccdc80 | protein_coding | -1.118728391 | 0.001008466 | 0.018635615 |
| ENSMUSG00000022817 | Itgb5 | protein_coding | -0.615658024 | 0.000239549 | 0.007129132 |
| ENSMUSG00000023961 | Enpp4 | protein_coding | 0.587731147 | 0.001786403 | 0.028232323 |
| ENSMUSG00000024122 | Pdpk1 | protein_coding | 0.541507352 | 0.000963049 | 0.018007376 |
| ENSMUSG00000024610 | Cd74 | protein_coding | -0.791664555 | 1.00556021315183e-06 | 0.000122361 |
| ENSMUSG00000024659 | Anxa1 | protein_coding | -1.512472665 | 4.25109248690384e-06 | 0.000384277 |
| ENSMUSG00000024696 | Lpxn | protein_coding | -0.872017849 | 0.001955225 | 0.03002613 |
| ENSMUSG00000024899 | Papss2 | protein_coding | 1.102825036 | 0.000307703 | 0.008518827 |
| ENSMUSG00000024909 | Efemp2 | protein_coding | -0.777232616 | 0.000700932 | 0.014643303 |
| ENSMUSG00000025225 | Nfkb2 | protein_coding | -1.088243654 | 0.000265238 | 0.007651077 |
| ENSMUSG00000025810 | Nrp1 | protein_coding | 0.657653065 | 0.001186178 | 0.020964182 |
| ENSMUSG00000026031 | Cflar | protein_coding | 0.532765574 | 0.000830436 | 0.016255934 |
| ENSMUSG00000026042 | Col5a2 | protein_coding | -1.206568275 | 0.000728819 | 0.014934605 |
| ENSMUSG00000026043 | Col3a1 | protein_coding | -1.440554499 | 7.85689078160858e-06 | 0.000592523 |
| ENSMUSG00000026131 | Dst | protein_coding | 0.666104263 | 7.67060124503912e-05 | 0.003155396 |
| ENSMUSG00000026177 | Slc11a1 | protein_coding | -0.971611166 | 8.98448038739113e-05 | 0.003571646 |
| ENSMUSG00000026193 | Fn1 | protein_coding | 0.596975959 | 0.003777928 | 0.045669807 |
| ENSMUSG00000026574 | Dpt | protein_coding | -2.276000376 | 2.43494983261836e-09 | 1.11473574272644e-06 |
| ENSMUSG00000026579 | F5 | protein_coding | 0.590975989 | 0.000147789 | 0.005078516 |
| ENSMUSG00000026728 | Vim | protein_coding | -1.008012231 | 0.000331247 | 0.008954399 |
| ENSMUSG00000027230 | Creb3l1 | protein_coding | -1.160455218 | 0.000277261 | 0.007822827 |
| ENSMUSG00000027447 | Cst3 | protein_coding | -0.599680104 | 0.000476341 | 0.01124831 |
| ENSMUSG00000027611 | Procr | protein_coding | -1.02543997 | 5.43884027460689e-05 | 0.002522484 |
| ENSMUSG00000027715 | Ccna2 | protein_coding | -1.785630846 | 2.12594381127603e-10 | 1.12082712802175e-07 |
| ENSMUSG00000027947 | Il6ra | protein_coding | 0.968885911 | 0.000261958 | 0.007628452 |
| ENSMUSG00000028284 | Map3k7 | protein_coding | 0.547480983 | 0.000892646 | 0.017089669 |
| ENSMUSG00000028339 | Col15a1 | protein_coding | -2.212241491 | 3.18249427641521e-09 | 1.36520115398135e-06 |
| ENSMUSG00000028369 | Svep1 | protein_coding | -1.393792067 | 0.002069319 | 0.031109932 |
| ENSMUSG00000028434 | Epb41l4b | protein_coding | 0.596587994 | 0.000188338 | 0.00603362 |
| ENSMUSG00000028469 | Npr2 | protein_coding | 0.909971092 | 0.000315828 | 0.008636292 |
| ENSMUSG00000028517 | Plpp3 | protein_coding | 0.805973297 | 6.50139016543545e-05 | 0.002831502 |
| ENSMUSG00000028583 | Pdpn | protein_coding | -2.134764817 | 0.002329449 | 0.033582114 |
| ENSMUSG00000028664 | Ephb2 | protein_coding | -2.07963198 | 5.22207694762235e-08 | 1.34748574619375e-05 |
| ENSMUSG00000029061 | Mmp23 | protein_coding | -0.848606891 | 0.003206267 | 0.041254163 |
| ENSMUSG00000029661 | Col1a2 | protein_coding | -1.337181846 | 2.36399793040986e-05 | 0.001380785 |
| ENSMUSG00000030342 | Cd9 | protein_coding | -1.173381532 | 0.000177081 | 0.005743216 |
| ENSMUSG00000030717 | Nupr1 | protein_coding | -2.637632114 | 1.9288682355559e-10 | 1.12082712802175e-07 |
| ENSMUSG00000031010 | Usp9x | protein_coding | 0.579073779 | 0.000201146 | 0.00629143 |
| ENSMUSG00000031196 | F8 | protein_coding | 0.607776218 | 0.000487037 | 0.011387204 |
| ENSMUSG00000031198 | Fundc2 | protein_coding | -0.688615518 | 0.000273867 | 0.007786415 |
| ENSMUSG00000031538 | Plat | protein_coding | -1.527781477 | 7.48602949659561e-05 | 0.003124757 |
| ENSMUSG00000031613 | Hpgd | protein_coding | 1.230658639 | 4.09714305893807e-11 | 2.90733271462245e-08 |
| ENSMUSG00000031778 | Cx3cl1 | protein_coding | -1.705710743 | 3.34259267575044e-05 | 0.001796897 |
| ENSMUSG00000031825 | Crispld2 | protein_coding | -1.554137049 | 0.002427958 | 0.034388804 |

| gene_id | external_gene_name | gene_biotype | log2FoldChange | pvalue | padj |
|---|---|---|---|---|---|
| ENSMUSG00000031838 | Ifi30 | protein_coding | -0.667019565 | 0.001476782 | 0.024459276 |
| ENSMUSG00000032020 | Ubash3b | protein_coding | -0.787052204 | 0.003600469 | 0.044336588 |
| ENSMUSG00000032334 | Loxl1 | protein_coding | -1.745265056 | 0.000260486 | 0.007606607 |
| ENSMUSG00000032431 | Crtap | protein_coding | -0.730881345 | 0.003829627 | 0.046059374 |
| ENSMUSG00000034205 | Loxl2 | protein_coding | -0.992895342 | 0.000134524 | 0.004725667 |
| ENSMUSG00000034652 | Cd300a | protein_coding | -1.019776784 | 1.48889776120713e-06 | 0.000163802 |
| ENSMUSG00000036545 | Adamts2 | protein_coding | -0.931160789 | 0.003114 | 0.040504656 |
| ENSMUSG00000036585 | Fgf1 | protein_coding | 1.972285711 | 6.08107376821681e-07 | 8.3788930988867e-05 |
| ENSMUSG00000036856 | Wnt4 | protein_coding | -1.302947492 | 0.000811084 | 0.016031892 |
| ENSMUSG00000036894 | Rap2b | protein_coding | -0.773677283 | 0.00114442 | 0.020481227 |
| ENSMUSG00000038642 | Ctss | protein_coding | -1.141910284 | 1.78918749435309e-06 | 0.000186707 |
| ENSMUSG00000038658 | Ric1 | protein_coding | 0.527042291 | 0.003408278 | 0.043034061 |
| ENSMUSG00000038745 | Nlrp6 | protein_coding | 0.602190164 | 0.002106936 | 0.031442309 |
| ENSMUSG00000039087 | Rreb1 | protein_coding | 0.633722089 | 0.001403358 | 0.023514125 |
| ENSMUSG00000039662 | Icmt | protein_coding | 0.560614133 | 0.000869224 | 0.016852494 |
| ENSMUSG00000041324 | Inhba | protein_coding | 1.368040414 | 0.000243564 | 0.007231494 |
| ENSMUSG00000041431 | Ccnb1 | protein_coding | -1.830365105 | 1.07458870743048e-06 | 0.000128156 |
| ENSMUSG00000042699 | Dhx9 | protein_coding | 0.482368196 | 0.003515081 | 0.043913761 |
| ENSMUSG00000043903 | Zfp469 | protein_coding | -2.151859219 | 0.002608717 | 0.036261428 |
| ENSMUSG00000046223 | Plaur | protein_coding | -1.336710244 | 2.14264555082897e-05 | 0.001299693 |
| ENSMUSG00000047407 | Tgif1 | protein_coding | -0.760380691 | 0.003291247 | 0.041891815 |
| ENSMUSG00000049723 | Mmp12 | protein_coding | -1.841464834 | 1.11018364721254e-07 | 2.39542899036847e-05 |
| ENSMUSG00000050335 | Lgals3 | protein_coding | -1.605813304 | 6.24088152263731e-07 | 8.48487616678074e-05 |
| ENSMUSG00000050989 | Selenon | protein_coding | -0.774054042 | 0.003842859 | 0.046179381 |
| ENSMUSG00000052384 | Nrros | protein_coding | -0.740538059 | 0.004230799 | 0.049237969 |
| ENSMUSG00000054342 | Kcnn4 | protein_coding | -1.935491933 | 8.59570392140514e-05 | 0.003440346 |
| ENSMUSG00000056071 | S100a9 | protein_coding | -1.706237111 | 0.001464127 | 0.0243028 |
| ENSMUSG00000058715 | Fcer1g | protein_coding | -1.08062574 | 3.21505730541532e-09 | 1.36520115398135e-06 |
| ENSMUSG00000059552 | Trp53 | protein_coding | -0.56833925 | 0.003534258 | 0.043996559 |
| ENSMUSG00000061436 | Hipk2 | protein_coding | 1.347467196 | 3.66520835620329e-14 | 5.20166369912371e-11 |
| ENSMUSG00000062960 | Kdr | protein_coding | 0.863015837 | 1.65107621920769e-07 | 3.27352200551177e-05 |
| ENSMUSG00000066113 | Adamtsl1 | protein_coding | -1.286770081 | 0.00224774 | 0.032717876 |
| ENSMUSG00000070436 | Serpinh1 | protein_coding | -1.017098162 | 0.000186917 | 0.006001658 |
| ENSMUSG00000079293 | Clec7a | protein_coding | -1.140493794 | 5.51659679318721e-07 | 7.75163779098147e-05 |

**Supplementary Table 3: Sets of comparisons and RNA-seq analysis in this study**

| Sets of analysis | Diet and treatment | Duration (weeks) | PCA | DEGs | GO enrichment analysis | KEGG analysis |
|---|---|---|---|---|---|---|
| CD vs HFD (Data from concurrent treatment file) | CD | 10 | / | Fig. 2a<br>Fig. 3b<br>Extended Data Fig. 2<br>Extended Data Fig. 6b | Fig. 1e (Lipofuscin)<br>Fig. 2b (Top 20)<br>Fig. 2c (Neutrophil)<br>Fig. 3a (Cytokine & chemokine) | Extended Data Fig. 6a |
| | HFD | 10 | | | | |
| CD vs HFD (Data from delayed treatment file) | CD | 17 | / | Fig. 6b | / | Fig. 6a |
| | HFD | 17 | | | | |
| CD vs HFD vs HFDT (Concurrent treatment) | CD | 10 | Extended Data Fig. 5a | Extended Data Fig. 5c-e<br>Extended Data Fig. 6c,d<br>Extended Data Fig. 7a | / | / |
| | HFD | 10 | | | | |
| | HFDT | 10 (HFD+TZP) | | | | |
| CD vs HFD vs HFDT (Delayed treatment) | CD | 17 | Fig. 5g | Fig. 5i-k<br>Fig. 6c,d<br>Extended Data Fig. 7b | / | / |
| | HFD | 17 | | | | |
| | HFDT | 7 (HFD) + 10 (HFD+TZP) | | | | |
| HFD vs HFDT (Concurrent treatment) | HFD | 10 | / | / | Extended Data Fig. 5b | / |
| | HFDT | 10 (HFD+TZP) | | | | |
| HFD vs HFDT (Delayed treatment) | HFD | 17 | / | / | Fig. 5h | / |
| | HFDT | 7 (HFD) + 10 (HFD+TZP) | | | | |
